# A Cell Viability and Utility Index (CVUI) for wildlife fibroblast biobanking: framework development and preliminary empirical validation

**DOI:** 10.64898/2026.07.31.742150

**Authors:** Natalie E. Calatayud, Leah Jacobs, Jennifer Hetz-Rodriguez, Yanni Chan, Kevin Rowe, Karen Rowe, Jane Melville, Karen Roberts, Kylea Clarke, Katie Date, Andrew J. Pask, Joanna Sumner

## Abstract

Wildlife biobanking depends on consistent, reproducible cell culture outcomes, yet no standardised framework exists for assessing culture quality across taxa or for integrating quality metrics into institutional collections management systems. Here, we present the Cell Viability and Utility Index (CVUI), a scored pipeline framework validated in a pilot study of 154 culture rounds from 46 species held in the Ian Potter Australian Wildlife Biobank at Museums Victoria Research Institute. The CVUI was adapted from the Wildlife Sperm Index (Jacobs et al., 2026) and assigns weighted scores encompassing four sequential culture stages — establishment, first passage, expansion, and cryobanking — with a continuous viability modifier applied at the cryobanking stage. Survival analysis identified individual animal identity as the primary source of variance in culture establishment, whereas taxon was the primary predictor of first-passage success. Beyond first passage, failure rates were too low to reliably assess either effect, with critical attrition concentrated at the earliest pipeline stages. These findings informed the differential weighting of CVUI components and have direct implications for workflow prioritisation and banking strategy. Monitoring can now be targeted at establishment and first passage, the stages where intervention has the greatest impact, with taxon-specific attention warranted at first passage and individual-level attention at establishment, while the robustness of the late-stage protocol means that banking yield is largely determined before a culture reaches P1. Integration of CVUI scores into collections management platforms such as Axiell EMu (Electronic Museum) would transform passive record-keeping into active decision support, enabling real-time quality tracking, early flagging of at-risk cultures, and longitudinal benchmarking of biobank performance across species, collectors, and tissue types. This framework provides a foundation for optimising wildlife fibroblast biobanking protocols, developing taxon-specific culture standards, and expanding standardised quality assessment across wildlife biobanks nationally and internationally.

## INTRODUCTION

Biodiversity loss is accelerating under the combined pressures of human-induced land use change, habitat fragmentation, and climate change. Genetic diversity, a critical Biodiversity Variable under the Aichi Targets and the Kunming-Montreal Global Biodiversity Framework (CBD, 2011; CBD, 2022), has declined an estimated six per cent since the Industrial Revolution, with losses most severe in fragmented and island populations (Leigh et al., 2019; Shaw et al., 2025). On-the-ground conservation efforts have proved insufficient given the scale of these losses (Burgin, 2008; McKenney & Kiesecker, 2010). Germplasm biobanks have emerged as a complement to conservation efforts, serving as long-term repositories for preserving genetic materials, including tissues, DNA, gametes, and living cells, to safeguard species diversity. These efforts support breeding and restoration programmes and provide a safety net for endangered taxa beyond the spatial and logistical limitations of captive management (Hvilsom et al., 2025; Mooney et al., 2023; Rola et al., 2025b). The number of cryopreservation studies published annually has increased significantly in recent decades, reflecting growing recognition of the conservation potential of biobanked material (Brereton et al., 2025). Somatic cells and stem cells have featured more frequently in the more recently published cryobanking literature, signalling a broader shift in the field toward cell types with greater long-term conservation utility (Brereton et al., 2025). Among cryopreserved materials, living somatic cell collections occupy a distinctive position, preserving not only genetic information but functional biological resources applicable to reproductive biotechnology, cytogenetics, and molecular research. Critically, somatic cells represent a foundational biological resource from which a broad range of approaches can be deployed, from induced pluripotent stem cell derivation and somatic cell nuclear transfer to genomic and transcriptomic profiling, offering pathways not only to study species at the cellular level but potentially to contribute to their recovery (Ben-Nun et al., 2011).

Primary fibroblast cultures derived from a range of somatic tissues, including skin, tongue, trachea, and other connective tissue sources, have become the dominant cell type used in wildlife biobanking, valued for their wide distribution throughout the body, relative ease of isolation, high replicative potential, and compatibility with a broad range of culture conditions across vertebrate taxa (Jiménez & Harper, 2023; Harper, 2024). Skin biopsies are particularly favoured in living animal collections as a minimally invasive source that does not require sedation or sacrifice, while other tissue types are routinely collected opportunistically from animals that have died in captivity or in the field. Despite this, establishing primary fibroblast cultures from wildlife poses considerable practical challenges. Culture success is influenced by a complex of interacting factors, including tissue source and biopsy site, post-mortem interval, transport conditions, and species-specific biological variation, with non-mammalian taxa in particular presenting technical difficulties that are not well characterised in the literature (Jiménez & Harper, 2023; Fernandes et al., 2016; Rola et al., 2025a). Outcomes vary not only between species but between individuals of the same species and across successive culture rounds from the same animal, reflecting the inherent biological variability of primary cell culture as a system.

The institutions best positioned to generate and curate living cell collections at scale — zoological institutions and natural history museums — have historically operated in relative isolation from one another despite holding deeply complementary resources. Zoos maintain longitudinal records on individual animals encompassing health, demography, pedigree, and tissue provenance, while natural history museums provide the curatorial infrastructure, long-term preservation capacity, and data management systems needed to make such material discoverable and comparable across institutions and taxa (Poo et al., 2022). Formal collaborations between these institution types remain surprisingly infrequent, and the scientific value of zoo-held biological material, including living cell collections, continues to be constrained by limited integration with broader biodiversity data infrastructure (Poo et al., 2022; Mooney et al., 2023). Addressing this gap requires not only institutional partnerships but also standardised, collections-compatible quality metrics that can make living cell data legible and actionable within the systems that natural history institutions already use.

Currently, standardised frameworks for assessing quality do not exist for culture quality and pipeline performance in wildlife fibroblast biobanking. Quality is typically reported as a binary outcome (e.g., passage number reached) or as post-thaw viability, with no agreed-upon metric that captures the full trajectory from tissue receipt to a cryobanked cell line, limiting comparability across institutions and taxa and preventing systematic identification of factors driving attrition (Wong et al., 2012). This gap mirrors a broader challenge identified across wildlife germplasm biobanking, where the absence of standardised quality metrics, inconsistent species representation, and a lack of shared data standards continue to constrain the utility and comparability of collections globally (Hvilsom et al., 2025; Mooney et al., 2023; Brereton et al., 2025). In established biobanking fields, standardised quality frameworks, encompassing tiered quality criteria, defined metadata requirements, and reproducible assessment protocols, are considered foundational to collection utility and downstream application (Rohani et al., 2018; Wong et al., 2012). The absence of equivalent standards in wildlife biobanking represents not merely an inconvenience but a fundamental barrier to realising the conservation potential of these collections.

Standardised composite indices have demonstrated considerable utility in addressing analogous gaps in quality assessment in wildlife reproductive biology. The Wildlife Sperm Index, developed to provide a single, weighted score capturing the multidimensional quality of wildlife spermatozoa across taxa, demonstrated that a weighted pipeline framework could substantially improve comparability and interpretive consistency across institutions and species (Jacobs et al., 2026). The Cell Viability and Utility Index, or CVUI, applies the same logic to primary fibroblast culture, providing a scored, weighted assessment of culture performance across the full pipeline from tissue preparation to cryobanking. Weights are derived empirically from observed attrition rates at each pipeline stage, focusing scoring on stages where biological failure is most likely to occur and ensuring the index reflects the true difficulty distribution of the culture process. Here we present the CVUI as a pilot framework developed and validated at the Ian Potter Australian Wildlife Biobank (IPAWB), Museums Victoria Research Institute, using a dataset of 154 culture rounds spanning 46 species across three taxonomic groups: birds, mammals, and reptiles. We describe the biological and statistical basis for the index, present a survival analysis identifying taxon as the primary predictor of first-passage failure, and demonstrate the application of the CVUI as a practical tool for quality benchmarking, workflow prioritisation, and longitudinal performance monitoring in wildlife fibroblast biobanking. The integration of CVUI scores into collections management platforms such as Axiell Electronic Museum (EMu), widely used across natural history museums, has the potential to transform static specimen records into active decision-support tools, enabling real-time quality tracking, longitudinal benchmarking across species and collections, and the identification of systemic workflow inefficiencies at an institutional scale. While demonstrated here within an EMu environment, the CVUI is designed as a platform-agnostic framework, readily adaptable to other collections management and laboratory information systems used by biobanking institutions worldwide.

## Methods

### Framework development

The CVUI was developed as a sequential, gated pipeline that reflects the discrete biological stages of primary fibroblast culture, from tissue preparation to cryobanking. Four outcomes were defined, each representing a critical decision point at which a culture either progresses or fails. Each outcome is gated on the success of the preceding stage — failure at any point terminates scoring for all subsequent outcomes. The dataset comprised all culture rounds initiated and completed at the IPAWB between 1 January and 31 December 2025; culture rounds still active at the time of data extraction fell outside this window by definition and were therefore not included (Figure 1).

**Figure 1.**
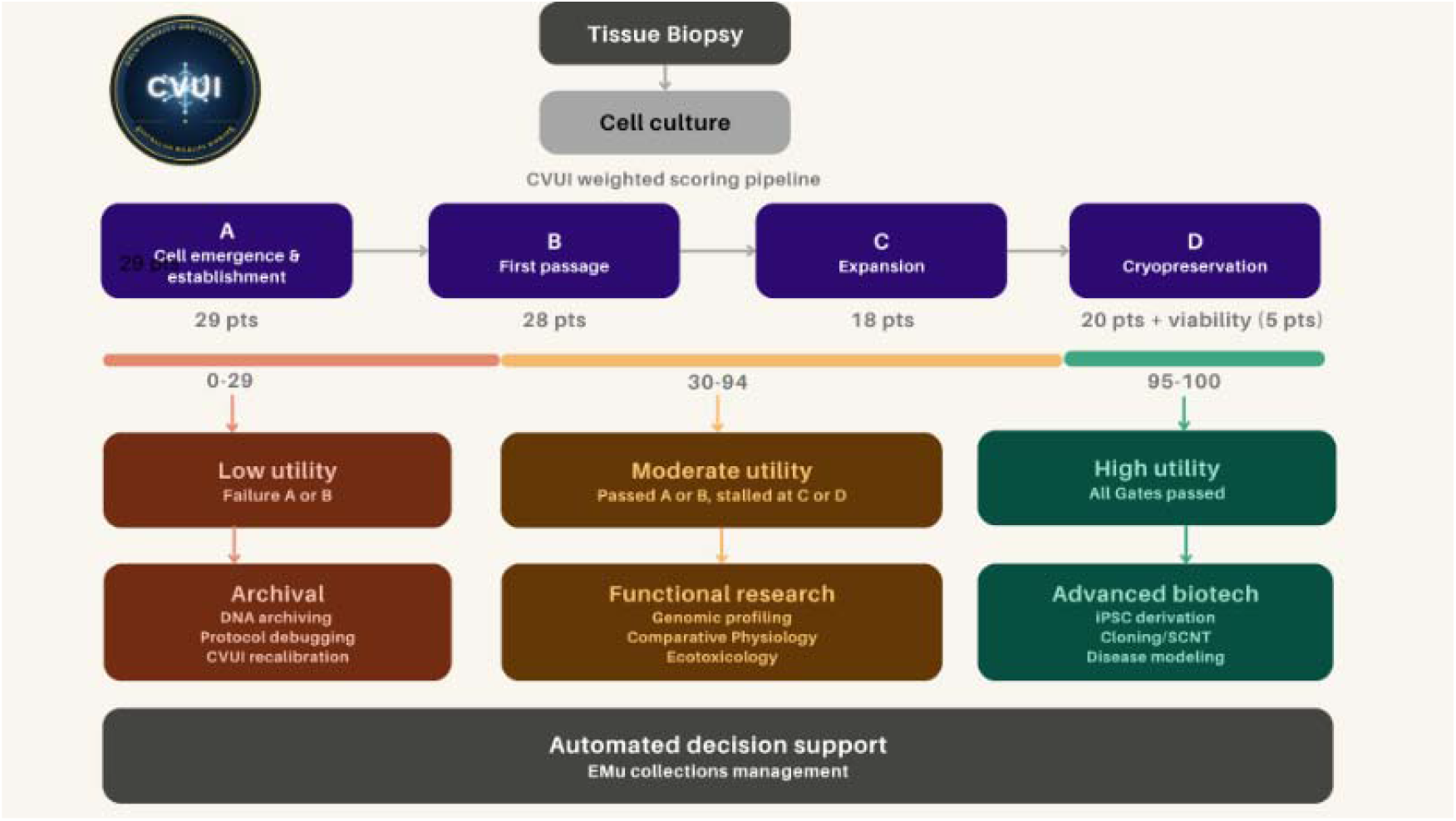
Cell Viability and Utility Index (CVUI) decision tree: illustrating the weighted scoring pipeline applied to wildlife somatic cell culture rounds following tissue biopsy and initiation of cell culture. Each round is evaluated against four sequential gates: A, cell emergence and establishment (29 points); B, first passage (28 points); C, expansion (18 points); and D, cryopreservation (20 points, plus up to 5 points for post-freeze viability). Points accumulate across gates successfully reached, producing a composite CVUI score out of 100. Scores fall into three utility tiers. Low utility (0–29) reflects failure at gate A or B and constrains a culture round to archival applications, including DNA archiving, protocol debugging, and CVUI recalibration. Moderate utility (30–94) reflects success at gate A or B but stalling at gate C or D, supporting functional research applications such as genomic profiling, comparative physiology, and ecotoxicology. High utility (95–100) reflects successful progression through all four gates, enabling advanced biotechnological applications such as iPSC derivation, cloning or somatic cell nuclear transfer (SCNT), and disease modelling. CVUI scores and associated metadata are intended for integration into EMu, Museums Victoria’s collection management system, to support automated decision-making for wildlife biobank holdings.

Time interval variables were derived from date-interval calculations between pipeline events: tissue preparation to first cell emergence, tissue preparation to discard, first cells to first passage, and first passage to second passage, each calculated in days from the recorded dates in the master dataset. A custom extraction pipeline was developed in R to apply consistent outcome definitions, derive these variables, and flag records with incomplete data. All extraction logic was version-controlled; the analyses presented here are based on Version 9 of the extraction script applied to the dataset described above.

### Preliminary empirical validation

The pilot dataset comprised 154 culture rounds from 141 unique dishes representing 46 species across three taxonomic groups: birds (n = 80 culture rounds), mammals (n = 60 culture rounds), and reptiles (n = 14 culture rounds); the difference between dish and round counts reflects dishes that were split or sub cultured into more than one round during the expansion process, such that a single original dish could give rise to multiple tracked culture rounds. Only rounds with a defined outcome were eligible for CVUI scoring at each stage; the number of eligible rounds differed by outcome due to the gated pipeline structure: 106 for Outcome A, 60 for Outcome B, 35 for Outcome C, and 32 for Outcome D (Table 1). Statistical modelling at each stage was further restricted to rounds with non-missing values for the relevant time-interval variable and, for the Cox mixed-effects models, for individual animal identity; the resulting analytic samples are reported below for each model.

**Table 1.** Outcome distribution across the culture pipeline by taxonomic group. Eligible rounds differ by outcome due to the gated pipeline structure. Success rates reflect the proportion of eligible rounds reaching each stage.

| Comparison | HR | 95% CI | p-value |
| --- | --- | --- | --- |
| Mammal vs Bird | 3.618 | 1.37–9.54 | 0.009 |
| Reptile vs Bird | 0.641 | 0.20–2.01 | 0.450 |
| Overall model ($\chi^2=8.35$, $df=3$) | | | 0.039 |
| Random effect SD (individual animal) | 0.020 |  |  |

To assess the relationship between time-interval variables and culture outcomes at each pipeline stage, univariate logistic regression was fitted separately for each outcome, with the relevant time-interval variable as the sole explanatory variable. Model discrimination was assessed using the area under the receiver operating characteristic curve, with values above 0.70 considered acceptable. To account for the non-independence of multiple culture rounds from the same individual animal, a Cox proportional hazards mixed effects model was fitted for each outcome using the coxme package in R, with individual animal identity as a random intercept and taxon as a fixed effect.

CVUI weights were derived empirically from observed stage-specific success rates in the pilot dataset, using each outcome’s success rate as an inverse weighting factor so that stages with higher attrition contributed proportionally more to the total score. For each outcome, an unscaled weight was calculated as the reciprocal of its observed success rate, and these reciprocals were scaled so that they summed to 95, reserving the remaining five points for the viability quality modifier described below. Formally, for outcome i:

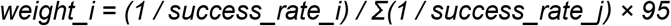

Observed success rates were 57% at Outcome A, 58% at Outcome B, 91% at Outcome C, and 81% at Outcome D. Applying this formula and rounding to the nearest integer using the largest-remainder method yielded weights of 29, 28, 18, and 20 for Outcomes A through D, respectively, summing to 95. A five-point quality modifier was applied to Outcome D, calculated as the proportion of live cells recorded at freeze, giving a maximum possible score of 100. Where a culture was successfully banked but viability data were absent, the modifier was set to one as a benefit of the doubt, yielding a score of 95 flagged for retrospective update. Where viability was recorded as zero with no cell concentration available, both the modifier and Outcome D score were set to zero. The final CVUI formula is:

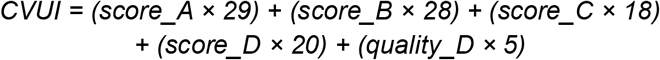

where score_A through score_D each take a value of one if the corresponding outcome was met and zero otherwise, and quality_D is a continuous variable bounded between zero and one. CVUI scores are the cumulative sum of weights for each gate successfully passed, not a percentage measure, and map onto discrete pipeline stages: zero indicates no cell establishment (no gates passed); 29 indicates establishment without progression to first passage (score_A only, 29); 57 indicates progression to first passage without expansion (score_A + score_B, 29 + 28); 75 indicates expansion without cryobanking (score_A + score_B + score_C, 29 + 28 + 18); and 95 to 100 indicates successful cryobanking with recorded viability (score_A + score_B + score_C + score_D, 29 + 28 + 18 + 20 = 95, plus 0 to 5 from the quality modifier). The weighting rationale and resulting score thresholds are illustrated in the CVUI decision tree (Figure 1).

## RESULTS

A total of 154 fibroblast cell culture rounds from 141 unique dishes representing 46 species across three taxonomic groups were included in the analysis. Culture rounds progressed through four sequential gated outcomes: establishment of first cells (Outcome A), progression to first passage (Outcome B), expansion to second passage or direct cryopreservation (Outcome C), and successful cryobanking (Outcome D). Outcome distribution and taxon-specific success rates are summarised in Table 1.

Overall establishment success at Outcome A was 56.6%, with Reptiles establishing at a markedly higher rate than Birds or Mammals. Success rates increased progressively across later pipeline stages, reaching 91% at Outcome C and 81% at Outcome D. The much larger absolute number of failures at Outcomes A and B (46 and 25, respectively) relative to Outcomes C and D (3 and 6) indicates that attrition is concentrated at the earliest pipeline stages, even though formal predictor testing was only reliably powered at Outcomes A and B. This pattern directly informed the differential weighting of CVUI components.

A Cox proportional hazards mixed effects model with taxon as a fixed effect and individual animal identity as a random intercept revealed a significant overall effect of taxon on progression to first passage (Outcome B; χ^2^=8.35, df=3, p=0.039; Table 3). Mammalian cultures had 3.6 times the hazard of failing to reach P1 compared to Bird cultures (HR=3.618, 95% CI: 1.37–9.54, p=0.009), while Reptile cultures did not differ significantly from Birds (HR=0.641, p=0.450). The random-effects variance at Outcome B was negligible (SD = 0.020), indicating that once taxon was accounted for, individual animal identity explained little additional variance in the first-passage outcome. This contrasts with Outcome A, where individual animal identity accounted for substantial variance in culture establishment (random-effects SD = 1.98), suggesting that animal-level factors, potentially including tissue quality, health status at collection, and individual variation in fibroblast proliferative capacity, are the primary determinants of whether a culture establishes.

At Outcomes C and D, the low absolute number of failures, three and six, respectively, reflects the high success rate of the protocol at later pipeline stages. Cox mixed-effects models were nonetheless fitted at both stages for consistency with Outcomes A and B; neither showed a significant effect of taxon (Outcome C: χ^2^=1.12, df=3, p=0.773;Outcome D: χ^2^=1.14, df=3, p=0.767), and the resulting hazard ratios were large in magnitude (Mammal HR=3.01 and 1.63; Reptile HR=3.81 and 2.46, at Outcomes C and D respectively) but not statistically distinguishable from no effect given the very small number of events. These results should be treated as inconclusive rather than as evidence of a negligible taxon effect at later pipeline stages and indicate that intervention efforts and predictive monitoring are most valuable and most reliably assessed at the establishment and first passage stages of the culture pipeline.

CVUI scores were calculated for all 106 eligible culture rounds, and summary statistics are presented in Table 2. The score distribution was markedly bimodal, with 46 rounds scoring zero and 26 scoring between 95 and 100, reflecting the gated nature of the pipeline in which cultures either fail to establish or once established, tend to progress through to cryobanking (Figure 2). Intermediate score categories were small, consistent with the high success rates at Outcomes C and D. The Mammal median of zero reflects the high proportion of rounds failing at establishment, while the Reptile mean of 50.1 reflects frequent establishment despite the smallest sample size. All 26 banked cultures had viability recorded, with no missing viability flags in the dataset. The continuous quality modifier is capped at 99.95, as no culture recorded exactly 100% viability at freeze. The distribution of CVUI scores across taxa and their relationship to pipeline stage thresholds are shown in Figure 3.

**Table 2.** CVUI score summary statistics. Overall and by taxonomic group (n=106 eligible culture rounds).

| Comparison | HR | 95% CI | p-value |
| --- | --- | --- | --- |
| Mammal vs Bird | 3.618 | 1.37–9.54 | 0.009 |
| Reptile vs Bird | 0.641 | 0.20–2.01 | 0.450 |
| Overall model ($\chi^2=8.35$, df=3) | | | 0.039 |
| Random effect SD (individual animal) | 0.020 |  |  |

**Table 3.**
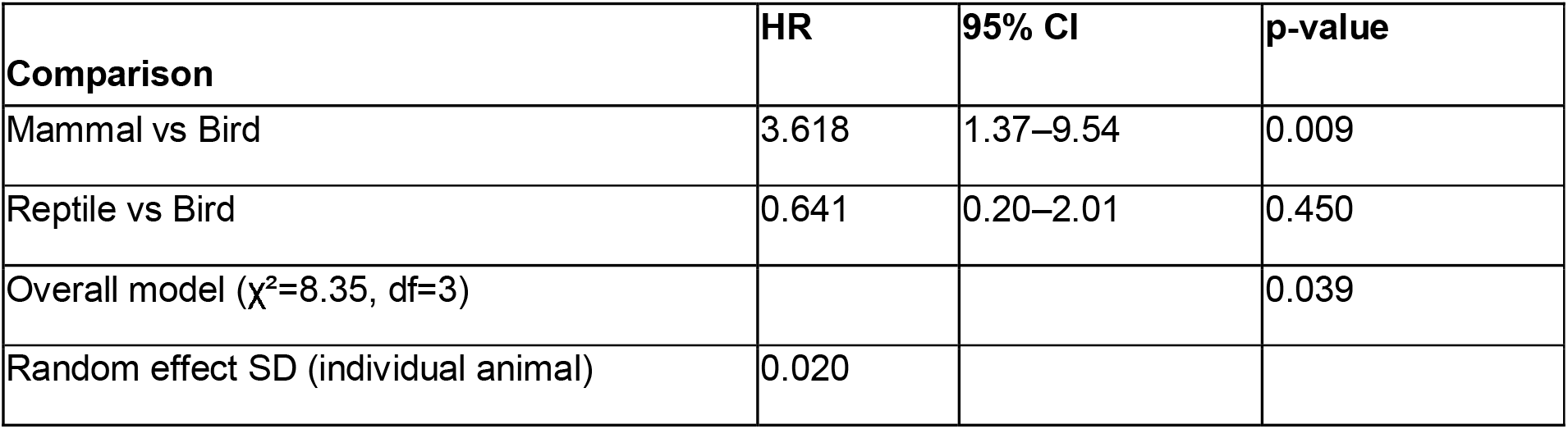
Cox proportional hazards mixed effects model results for Outcome B (progression to first passage). Individual animal identity was included as a random intercept. Bird is the reference taxon. HR = hazard ratio, CI = confidence interval.

**Figure 2.**
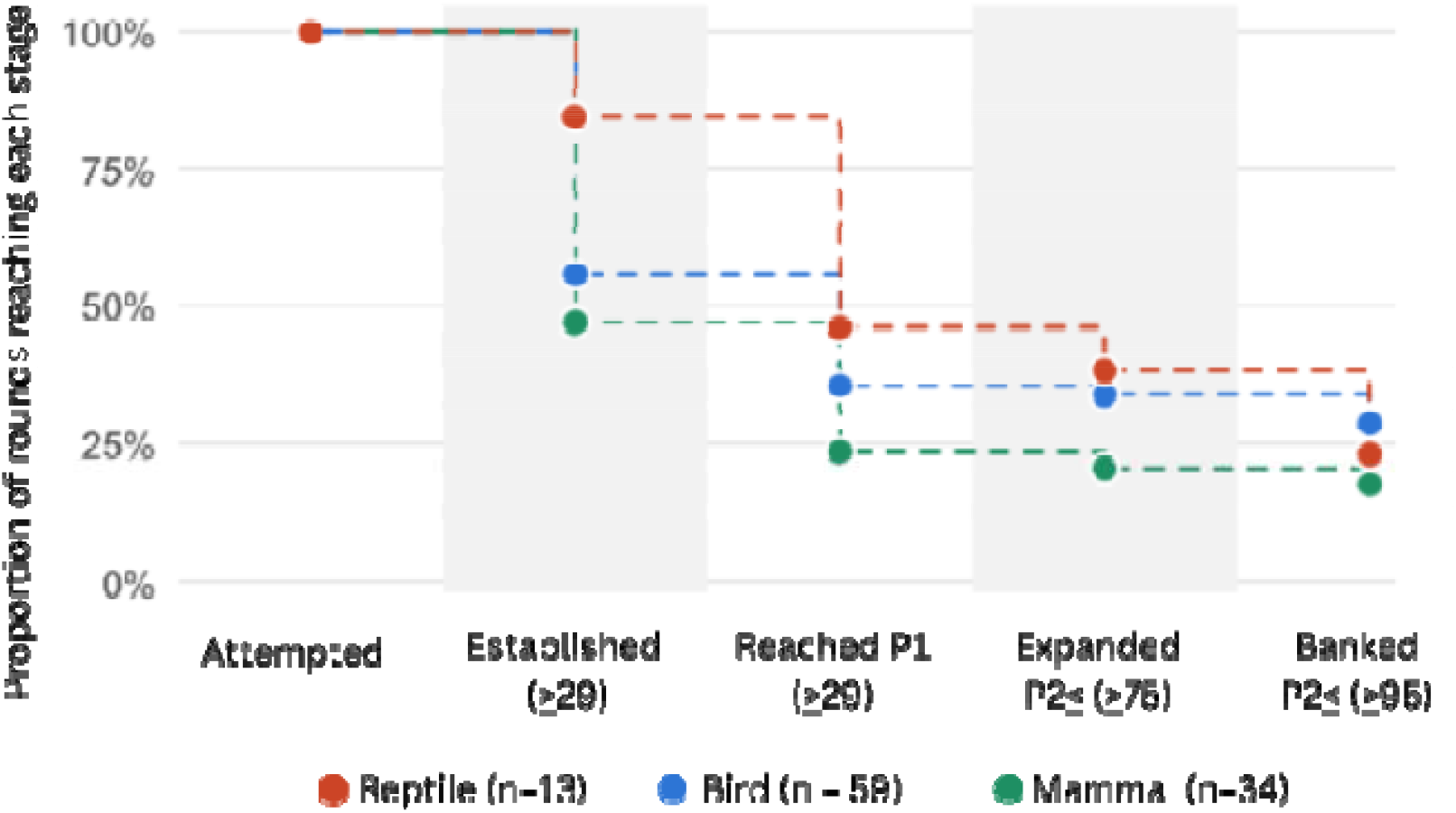
CVUI score category distribution by taxonomic group (n=106 eligible culture rounds). Cumulative attrition of culture rounds through the CVUI pipeline, by taxonomic group (n=106). Lines show the proportion of each taxon’s rounds reaching at least each stage: established (≥29), reached P1 (≥57), expanded (≥75), banked (≥95). Reptiles show the steepest early attrition; all three taxa converge toward similar banking rates (Bird n=59, Mammal n=34, Reptile n=13).

**Figure 3.**
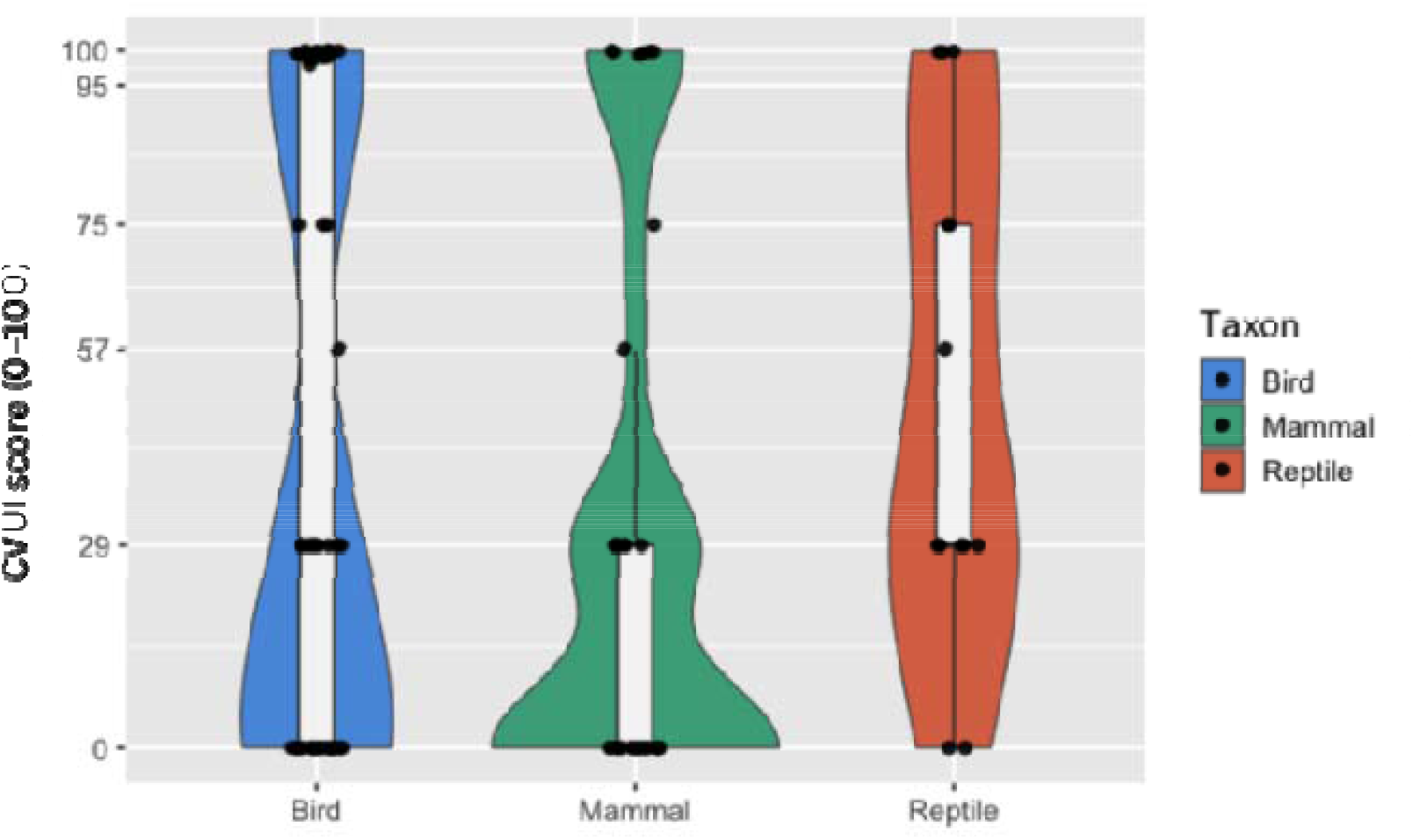
Distribution of Cell Viability and Utility Index (CVUI) scores by taxonomic group (n=106 culture rounds). Coloured violins show the density of scores within each taxon; the white box shows the median and interquartile range; black points show individual culture rounds, jittered horizontally for visibility. The bimodal shape reflects the gated pipeline structure, with cultures either failing to establish (score = 0) or progressing to cryobanking (score ≥ 95). Mammalian cultures had the lowest median CVUI score, consistent with a significantly elevated hazard of first-passage failure compared with Birds (HR = 3.618, p = 0.009). Bird n=59, Mammal n=34, Reptile n=13.

## DISCUSSION

The most substantive empirical finding of this study is the significantly elevated hazard of first passage failure in mammalian cultures relative to Birds, with Mammals showing 3.6 times the hazard of failing to reach P1 after accounting for individual animal identity. This finding is consistent with the known biological diversity of the Mammal group, which in this dataset spans a broad range of orders, body sizes, and ecological contexts. This result likely reflects genuine interspecific variation in fibroblast proliferative capacity and sensitivity to culture conditions (Jiménez & Harper, 2023; Harper, 2024), as well as variation in tissue quality at the time of collection (Fernandes et al., 2016; Rola et al., 2025a), rather than a systematic protocol failure. Mammals were not, however, the largest taxonomic group in this dataset: Birds comprised 80 of the 154 culture rounds, compared with 60 for Mammals and 14 for Reptiles, and, specifically at Outcome B, more Bird rounds were eligible than Mammal rounds (33 versus 16). The elevated Mammal hazard, therefore, cannot be attributed to a larger or more heterogeneous Mammal sample driving statistical power to detect attrition; rather, the taxonomic breadth noted above may itself be a genuine contributor to the elevated and variable hazard observed. A further unmeasured source of heterogeneity is sample provenance: tissue collected from captive animals during routine veterinary procedures is likely to differ systematically in baseline quality from tissue collected from deceased wild animals with variable post-mortem intervals (Rola et al., 2025a) or from roadkill specimens, reflecting differences in tissue type, body region, and condition at collection (Fernandes et al., 2016). Provenance was not consistently recorded in the current dataset and could not be included as a covariate. Disentangling taxon-level biological variation from provenance-related tissue quality is an important direction for future CVUI iterations as these metadata fields are captured more systematically.

The substantial individual-animal variance in culture establishment (Outcome A) is a biologically meaningful finding that extends beyond its statistical role as a model correction. Once the taxon is accounted for, a large proportion of variance in whether a culture establishes at all is attributable to which individual animal the tissue came from rather than to any measured predictor. This is consistent with what practitioners observe in the field — some animals consistently yield tractable cultures while others do not, regardless of species or protocol. Animal-level factors likely contributing to this variance include health status and condition at the time of collection, post-mortem interval where tissue was collected from deceased animals (Rola et al., 2025a), tissue type and body region (Fernandes et al., 2016), and intrinsic biological variation in cell viability between individuals of the same species (Jiménez & Harper, 2023). Post-mortem interval is a particularly plausible candidate given its established effect on tissue and cell viability, but this variable was recorded for only 14.9% of culture rounds in the current dataset, precluding its inclusion as a covariate in the present analysis. The current CVUI does not explicitly capture these variables, and incorporating animal-level covariates as additional index components is a clear priority for future versions.

This study establishes a foundation for several important future directions. The pilot dataset is drawn from a single year’s results, and while 46 species across three taxa provide a meaningful biological range, expanding the dataset across multi-year data aggregation and potentially across additional institutions will be essential for supporting taxon-level modelling at finer resolution and robustly characterising outcome patterns across a broader range of species. Amphibians were excluded from this analysis due to the well-documented technical challenges of establishing fibroblast cultures from this group (Jiménez & Harper, 2023; Zimkus et al., 2018), and extending the CVUI to amphibian cultures represents a clear and worthwhile future direction given the conservation urgency facing this taxon globally, and the growing recognition that standardised, integrative frameworks for amphibian genetic resource preservation are essential to realising the conservation potential of biobanked material (Zimkus et al., 2018).

The current version of the CVUI evaluates a single binary outcome at each pipeline stage, recording whether a culture progressed or failed without capturing the specific mode of failure. A valuable next step would be to layer stage-specific quality diagnostics onto the existing framework, for example, distinguishing failure due to insufficient proliferation, bacterial or fungal contamination, or low post-thaw viability, which would allow the index to move from documenting that attrition occurred to explaining why, sharpening both troubleshooting at the bench and the biological interpretation of the taxon- and individual-level effects reported here. Relatedly, incorporating finer-grained timing data — such as the interval between animal collection or death and tissue plating, and from collection or death to first cell emergence — would further extend the index’s diagnostic resolution upstream of the culture stages currently captured. The precision and interpretability of such timing data would be substantially improved by systematically recording sample provenance, for example, distinguishing tissue sourced from roadkill, euthanasia, or opportunistic wild collection from tissue obtained through captive collection, since time of death, and therefore post-mortem interval, can be defined far more narrowly for euthanised or captive-sourced material than for roadkill or opportunistically collected wild specimens, where time of death is frequently estimated rather than precisely known. Completeness of time-interval data was also limited at several pipeline stages, particularly for discard dates, and prospective standardisation of data collection will unlock the full power of survival analyses at Outcome A and improve the precision of future index iterations. This is particularly relevant in the context of frozen collections metadata, which, as a field, is still relatively young. Standardised frameworks for data collection in wildlife biobanking have been actively developed and refined only over the past five years or so, and it is well recognised that establishing consistent, institution-wide data standards takes considerable time and iterative refinement. These candidate pre-culture determinants, together with the post-thaw outcomes discussed below as a proposed fifth pipeline stage, are summarised alongside the outcomes tested in the current study in Figure 4.

**Figure 4.**
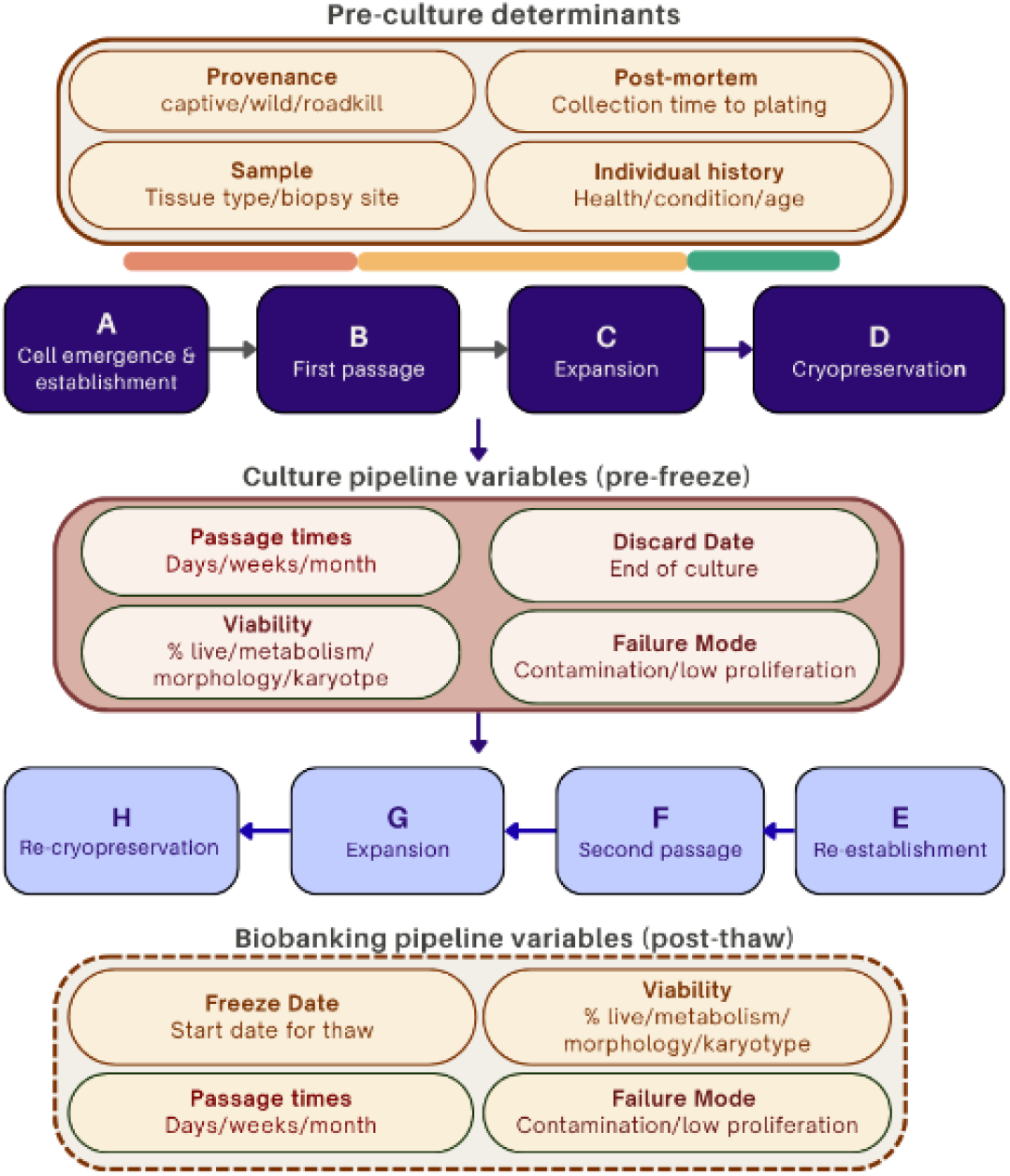
Determinants and future outcome tiers surrounding the tested CVUI pipeline. Determinants and future outcome tiers surrounding the tested CVUI pipeline. The four outcomes evaluated in this study (Outcome A, cell emergence and establishment; Outcome B, first passage; Outcome C, expansion; Outcome D, cryobanking) are shown centrally. Pre-culture determinants proposed as candidate covariates for future CVUI iterations are shown above. Culture pipeline variables recorded across the pre-freeze stages (passage timing, viability, discard date, and failure mode) are shown below Outcomes A–D. A proposed post-thaw outcome tier (Outcome E, re-establishment; Outcome F, second passage; Outcome G, expansion; Outcome H, re-cryopreservation) mirrors this same gated logic in reverse, reflecting the sequence in which a thawed culture would re-traverse the pipeline. Biobanking pipeline variables tracked at this post-thaw stage (freeze date, viability, passage timing, and failure mode) are shown at the base of the figure, allowing direct comparison with their pre-freeze equivalents above.

The CVUI is well-suited to this evolving landscape precisely because of its adaptive management capabilities: as weights are derived empirically from accumulating data, the index can be updated and recalibrated as metadata quality improves, new pipeline stages are added, and broader taxonomic and institutional representation is achieved. Rather than requiring a fully mature data infrastructure before it can be applied, the CVUI is designed to grow alongside the collections it monitors. As multi-year, multi-institutional datasets accumulate, the framework also provides a foundation for formal predictive modelling of culture success from pre-culture determinants — including post-mortem interval, tissue type, taxon, provenance, and collector identity — potentially enabling prospective triage of incoming samples before culture is initiated and shifting the index from a retrospective scoring tool to a prospective decision-support system. The need for greater collaboration between cryobanks and the adoption of shared data standards has been identified as one of the most pressing priorities in the field, with many institutions currently operating in isolation and the contents of individual cryobanks remaining largely unknown to other facilities (Brereton et al., 2025). The CVUI addresses this directly by providing a common quality language that can be adopted across institutions, enabling meaningful cross-institutional benchmarking and reducing the fragmentation that currently limits the collective conservation impact of living cell collections. Efforts to standardise biological material collection and quality assessment in vertebrate research have highlighted the importance of consistent metadata, provenance documentation, and tiered quality criteria as foundations for collections that can support future research and conservation applications (Wong et al., 2012), and the CVUI builds directly on this principle by providing a framework within which such standardisation can be systematically pursued and measured over time.

The global cryobanking community has identified the absence of standardised quality metrics and inconsistent species representation as key barriers to realising the conservation potential of living cell collections (Mooney et al., 2023; Hvilsom et al., 2025). A systematic review of the cryobanking literature confirms that the number of publications on wildlife cryopreservation priorities is significantly growing year on year, yet no consensus exists on the quality criteria, sample numbers, or collection standards that should guide biobanking decisions (Brereton et al., 2025). Many institutions continue to operate according to institution-specific priorities that limit cross-institutional comparability and knowledge sharing, and the CVUI represents a direct response to this gap at the level of individual culture performance. By translating culture outcomes into a single composite score empirically derived from observed attrition rates, the CVUI captures the full trajectory from tissue preparation to cryobanking in a form directly comparable across species, institutions, and time, extending the logic of the Wildlife Sperm Index (Jacobs et al., 2026) to somatic cell culture for the first time and converting routine culture records into actionable quality information without requiring additional assays or infrastructure beyond what is already collected in standard biobanking practice. The development of quality standards in biobanking follows a recognised trajectory across fields, from ad hoc institutional practice toward formalised, reproducible frameworks — and wildlife biobanking has now reached the point where this transition is both possible and necessary (Rohani et al., 2018).

Realising that potential at scale, however, depends equally on how quality data are stored, shared, and made discoverable across institutions. Zoos and natural history museums together hold the richest repositories of biological material from wildlife, yet the data systems underpinning these collections have historically been poorly integrated, limiting the scientific value of holdings that are, in many cases, irreplaceable (Poo et al., 2022). The integration of CVUI scoring into collections management platforms such as EMu — a system widely used across natural history collections for documentation, data management, and web export (Sendino, 2009) — is a direct step toward bridging this divide, enabling real-time score calculation, longitudinal benchmarking across species and collections, and the systematic identification of workflow inefficiencies at an institutional scale. Critically, making CVUI scores visible within the same infrastructure that curates specimen provenance, taxonomy, and tissue metadata positions cell quality data within the broader extended specimen framework, where a single animal’s biological record can be interrogated across its living history, its banked material, and its preserved voucher (Poo et al., 2022). While demonstrated here within a Museums Victoria context, the framework is designed to be platform-agnostic and readily adaptable to other laboratory information and collections management systems used by biobanking institutions worldwide.

The CVUI is presented here as a pilot framework, and its weights and thresholds should be treated as empirically derived starting points subject to revision as data accumulate across institutions and taxa. The current weighting scheme is deliberately simple, calculated directly from observed success rates rather than fitted statistically; as the dataset grows in scale and taxonomic breadth, there is clear scope to move toward weights, or even individual outcome predictions, derived directly from the survival and regression models presented here, potentially evolving the CVUI from a heuristic scoring tool into a statistically modelled, and eventually computational, framework for predicting culture outcomes from biological and procedural covariates. The broader value of high-quality, well-characterised living cell collections is increasingly evident — cryopreserved fibroblasts have already been used to generate induced pluripotent stem cells from nearly extinct species, opening pathways to assisted reproduction and potential genetic rescue that were previously impossible (Ben-Nun et al., 2011; Harper, 2024). Realising this potential depends not only on collecting cells but on characterising their quality and on having tools to track and systematically improve collection quality over time — tools that, as outlined above, could themselves grow more statistically sophisticated as the underlying dataset matures. Post-thaw recovery data are currently being generated from equivalent tissues and cells frozen in parallel, corresponding to the same individuals and tissue types used in this initial analysis. These data will inform future iterations of the index, with the goal of incorporating post-thaw re-establishment as an additional weighted pipeline stage, and represent a concrete, near-term opportunity to trial weights derived from fitted statistical models rather than from observed success rates alone. The CVUI provides that foundation. As wildlife fibroblast biobanking expands in scale and application, standardised quality metrics will be essential for ensuring that living cell collections deliver on their conservation potential.

## Supporting information

CVUI R-script

