## Supplementary material for "A Cell Viability and Utility Index (CVUI) for wildlife fibroblast biobanking: framework development and preliminary empirical validation": CVUI R-script

Supplementary Code S1

*CVUI survival analysis and data-processing pipeline (R Markdown, Version 11)*

This supplementary file provides the complete R Markdown script used to process raw culture-round records, construct the Cell Viability and Utility Index (CVUI), and perform the mixed-effects Cox survival analyses reported in the accompanying manuscript. The script is reproduced in full below, unmodified, as run to generate the results and figures referenced in the main text.

````
---
title: "CVUI as a Survival Analysis"
output: html_notebook
---

This is an [R Markdown](http://rmarkdown.rstudio.com) Notebook. When you execute code within the notebook, the results appear beneath the code.

Try executing this chunk by clicking the *Run* button within the chunk or by placing your cursor inside it and pressing *Cmd+Shift+Enter*.


#CVUI — Cell culture data: Outcome A to D
```{r}
# ============================================================
### CVUI — Cell culture data: Outcome A to D
### Version 9 — discard_date added (col 42)
# ============================================================

library(readr)
library(dplyr)
library(tidyr)
library(stringr)

# ------------------------------------------------------------
### STEP 1 — Load raw CSV
# ------------------------------------------------------------

raw <- read_csv(
  "/Users/nataliecalatayud/Documents/STATS & DATA FILES/R-STUDIO/CVUI/Cells - 2025_V2_Rstudio.csv",
  col_types      = cols(.default = "c"),
  name_repair    = "minimal",
  show_col_types = FALSE,
  locale         = locale(encoding = "latin1")
)

cat("Raw dimensions:", nrow(raw), "rows x", ncol(raw), "cols\n")

# ------------------------------------------------------------
### STEP 2 — Select and rename columns
# ------------------------------------------------------------

cells_raw <- raw %>%
  select(
    entry_nature   = 3,
    mv_parent      = 5,
    dish_id        = 6,
    taxa           = 18,
    common_name    = 19,
    species        = 20,
    date_collected = 16,
    date_prep      = 33,
    discard_date   = 42,    # NEW — was free text "Discarded/notes"
    date_1st       = 43,
    tissue_type    = 29,
    dish_size      = 45,
    p1             = 46,
    p2             = 47,
    p3             = 48,
    p4             = 49,
    p5             = 50,
    freeze         = 51,
    reg_no         = 52,
    cell_raw       = 53,
    live_raw       = 54,
    notes          = 55
  )

cat("Columns selected:", ncol(cells_raw), "\n")

# ------------------------------------------------------------
### STEP 3 — Filter to taxa of interest
# ------------------------------------------------------------

taxa_keep <- c("Bird", "Mammal", "Reptile")

cells_raw <- cells_raw %>%
  mutate(taxa = str_trim(taxa)) %>%
  filter(taxa %in% taxa_keep)

cat("Rows after taxa filter:", nrow(cells_raw), "\n")

# ------------------------------------------------------------
### STEP 4 — Parse dates
# ------------------------------------------------------------

parse_date_long <- function(x) {
  x <- str_trim(x)
  x[is.na(x) | x == "" | x == "NA" | x == "lost"] <- NA
  x[str_detect(x, regex(
    "discard|contam|failed|ongoing|no growth|not grow",
    ignore_case = TRUE
  ))] <- NA
  suppressWarnings(as.Date(x, format = "%d %B %Y"))
}

cells_raw <- cells_raw %>%
  mutate(across(
    c(date_collected, date_prep, discard_date, date_1st,
      p1, p2, p3, p4, p5, freeze),
    parse_date_long
  ))

# ------------------------------------------------------------
### STEP 5 — Parse cell_n
# ------------------------------------------------------------

parse_cell_n <- function(x) {
  x <- str_trim(x)
  x[is.na(x) | x == "" | x == "NA"] <- NA

  sapply(x, function(val) {
    if (is.na(val)) return(NA_real_)
    m6 <- regmatches(val, regexpr(
      "[0-9]+\\.?[0-9]*\\s*[xX]\\s*10\\^?6", val, perl = TRUE))
    m5 <- regmatches(val, regexpr(
      "[0-9]+\\.?[0-9]*\\s*[xX]\\s*10\\^?5", val, perl = TRUE))
    m4 <- regmatches(val, regexpr(
      "[0-9]+\\.?[0-9]*\\s*[xX]\\s*10\\^?4", val, perl = TRUE))
    if (length(m6) > 0) {
      return(as.numeric(str_extract(m6[1], "[0-9]+\\.?[0-9]*")) * 1e6)
    } else if (length(m5) > 0) {
      return(as.numeric(str_extract(m5[1], "[0-9]+\\.?[0-9]*")) * 1e5)
    } else if (length(m4) > 0) {
      return(as.numeric(str_extract(m4[1], "[0-9]+\\.?[0-9]*")) * 1e4)
    }
    return(NA_real_)
  }, USE.NAMES = FALSE)
}

cells_raw <- cells_raw %>%
  mutate(cell_n = parse_cell_n(cell_raw))

# ------------------------------------------------------------
### STEP 6 — Parse pct_live
# ------------------------------------------------------------

parse_live_pct <- function(x) {
  x <- str_trim(x)
  x[is.na(x) | x == "" | x == "?" | x == "#VALUE!"] <- NA

  sapply(x, function(val) {
    if (is.na(val)) return(NA_real_)
    val <- str_remove_all(val, "%")
    nums <- suppressWarnings(
      as.numeric(str_extract_all(val, "[0-9]+\\.?[0-9]*")[[1]])
    )
    nums <- nums[!is.na(nums) & nums >= 1 & nums <= 100]
    if (length(nums) == 0) return(NA_real_)
    mean(nums)
  }, USE.NAMES = FALSE)
}

cells_raw <- cells_raw %>%
  mutate(pct_live = parse_live_pct(live_raw))

# ------------------------------------------------------------
### STEP 7 — Classify entry_nature into round_status
# ------------------------------------------------------------

cells_raw <- cells_raw %>%
  mutate(
    entry_clean  = str_trim(str_to_lower(entry_nature)),
    round_status = case_when(

      # EXCLUDE — not culture attempts
      str_detect(entry_clean, "tissue source")    ~ "exclude",
      str_detect(entry_clean, "dna sample")       ~ "exclude",
      str_detect(entry_clean, "vitrification")    ~ "exclude",
      str_detect(entry_clean, "frozen cells")     ~ "exclude",
      str_detect(entry_clean, "slow freeze")      ~ "exclude",

      # CULTURE ROUNDS
      str_detect(entry_clean, "ongoing")          ~ "active",
      str_detect(entry_clean, "discarded")        ~ "discarded",
      str_detect(entry_clean, "done")             ~ "complete",
      str_detect(entry_clean, "finished")         ~ "complete",
      str_detect(entry_clean, "successful")       ~ "complete",
      str_detect(entry_clean, "almost done")      ~ "complete",
      str_detect(entry_clean, "same animal done") ~ "complete",
      str_detect(entry_clean, "culture ongoing")  ~ "active",
      str_detect(entry_clean, "culture finished") ~ "complete",
      str_detect(entry_clean, "ongoing plate")    ~ "active",
      str_detect(entry_clean, "cells ongoing")    ~ "active",
      is.na(entry_clean)                          ~ NA_character_,
      TRUE                                        ~ NA_character_
    )
  )

cat("Entry_nature classification check:\n")
cells_raw %>% count(round_status, sort = TRUE) %>% print(n = Inf)

# ------------------------------------------------------------
### STEP 8 — Remove excluded rows
# ------------------------------------------------------------

cells_raw <- cells_raw %>%
  filter(round_status != "exclude" | is.na(round_status))

cat("Rows after excluding tissue banking records:", nrow(cells_raw), "\n")

# ------------------------------------------------------------
### STEP 9 — Aggregate to dish × round level
# ------------------------------------------------------------

cells_rounds <- cells_raw %>%
  mutate(
    round_key = paste0(dish_id, "||", as.character(p1))
  ) %>%
  group_by(dish_id, round_key) %>%
  summarise(
    taxa           = first(na.omit(taxa)),
    mv_parent      = first(na.omit(mv_parent)),
    common_name    = first(na.omit(common_name)),
    species        = first(na.omit(species)),
    tissue_type    = first(na.omit(tissue_type)),
    dish_size      = first(na.omit(dish_size)),
    entry_nature   = first(na.omit(entry_nature)),
    entry_clean    = first(na.omit(entry_clean)),
    round_status   = first(na.omit(round_status)),
    date_collected = first(na.omit(date_collected)),
    date_prep      = first(na.omit(date_prep)),
    discard_date   = first(na.omit(discard_date)),
    date_1st       = first(na.omit(date_1st)),
    p1_date        = first(na.omit(p1)),
    p2_date        = first(na.omit(p2)),
    p3_date        = first(na.omit(p3)),
    p4_date        = first(na.omit(p4)),
    p5_date        = first(na.omit(p5)),
    freeze_date    = first(na.omit(freeze)),
    cell_n         = first(na.omit(cell_n)),
    pct_live       = first(na.omit(pct_live)),
    tube_n         = sum(!is.na(reg_no) & reg_no != ""),
    .groups = "drop"
  )

cat("Rows after aggregation to dish x round:", nrow(cells_rounds), "\n")

# ------------------------------------------------------------
### STEP 10 — Assign round number per dish
# ------------------------------------------------------------

cells_rounds <- cells_rounds %>%
  group_by(dish_id) %>%
  arrange(p1_date, .by_group = TRUE) %>%
  mutate(
    round_number  = row_number(),
    n_rounds_dish = n()
  ) %>%
  ungroup()

# ------------------------------------------------------------
### STEP 11 — Derive time intervals
# ------------------------------------------------------------

cells_rounds <- cells_rounds %>%
  mutate(
    days_collection_to_prep = as.numeric(date_prep   - date_collected),
    days_prep_to_first      = as.numeric(date_1st    - date_prep),
    days_first_to_p1        = as.numeric(p1_date     - date_1st),
    days_p1_to_p2           = as.numeric(p2_date     - p1_date),
    days_p2_to_freeze       = as.numeric(freeze_date - p2_date),
    days_prep_to_freeze     = as.numeric(freeze_date - date_prep),
    # NEW — time intervals using discard_date
    days_prep_to_discard    = as.numeric(discard_date - date_prep),
    days_first_to_discard   = as.numeric(discard_date - date_1st),
    days_p1_to_discard      = as.numeric(discard_date - p1_date),
    across(starts_with("days_"), ~ if_else(.x < 0, NA_real_, .x)),
    days_p1_to_p2           = if_else(days_p1_to_p2      > 200, NA_real_, days_p1_to_p2),
    days_p2_to_freeze       = if_else(days_p2_to_freeze   > 200, NA_real_, days_p2_to_freeze),
    days_first_to_p1        = if_else(days_first_to_p1    > 200, NA_real_, days_first_to_p1),
    days_prep_to_first      = if_else(days_prep_to_first  > 200, NA_real_, days_prep_to_first),
    days_collection_to_prep = if_else(days_collection_to_prep > 365, NA_real_, days_collection_to_prep)
  )

# ------------------------------------------------------------
### STEP 12 — Max passage reached
# ------------------------------------------------------------

cells_rounds <- cells_rounds %>%
  mutate(
    max_passage_reached = case_when(
      !is.na(p5_date) ~ 5L,
      !is.na(p4_date) ~ 4L,
      !is.na(p3_date) ~ 3L,
      !is.na(p2_date) ~ 2L,
      !is.na(p1_date) ~ 1L,
      TRUE            ~ 0L
    ),
    max_passage_label = case_when(
      max_passage_reached == 5L ~ "P5",
      max_passage_reached == 4L ~ "P4",
      max_passage_reached == 3L ~ "P3",
      max_passage_reached == 2L ~ "P2",
      max_passage_reached == 1L ~ "P1",
      TRUE                      ~ "No passage"
    )
  )

# ------------------------------------------------------------
### STEP 13 — Define gated outcomes A, B, C, D
# ------------------------------------------------------------

cells_analysis <- cells_rounds %>%
  mutate(

    # Outcome A — Plating to first cells
    outcome_A = case_when(
      is.na(date_prep)                               ~ NA_integer_,
      round_status == "active"                       ~ NA_integer_,
      is.na(round_status)                            ~ NA_integer_,
      !is.na(date_1st)                               ~ 1L,
      is.na(date_1st) & round_status %in%
        c("complete", "discarded")                   ~ 0L,
      TRUE                                           ~ NA_integer_
    ),

    # Outcome B — First cells to P1
    outcome_B = case_when(
      is.na(outcome_A) | outcome_A == 0L             ~ NA_integer_,
      round_status == "active"                       ~ NA_integer_,
      !is.na(p1_date)                                ~ 1L,
      is.na(p1_date) & round_status %in%
        c("complete", "discarded")                   ~ 0L,
      TRUE                                           ~ NA_integer_
    ),

    # Outcome C — P1 to expansion
    outcome_C = case_when(
      is.na(outcome_B) | outcome_B == 0L             ~ NA_integer_,
      round_status == "active"                       ~ NA_integer_,
      !is.na(p2_date)                                ~ 1L,
      is.na(p2_date) & !is.na(freeze_date)           ~ 1L,
      is.na(p2_date) & is.na(freeze_date) &
        round_status %in%
        c("complete", "discarded")                   ~ 0L,
      TRUE                                           ~ NA_integer_
    ),

    # Outcome D — Cryobanking
    outcome_D = case_when(
      is.na(outcome_C) | outcome_C == 0L             ~ NA_integer_,
      round_status == "active"                       ~ NA_integer_,
      !is.na(freeze_date)                            ~ 1L,
      is.na(freeze_date) & round_status %in%
        c("complete", "discarded")                   ~ 0L,
      TRUE                                           ~ NA_integer_
    )
  )

# ------------------------------------------------------------
### STEP 14 — Classify failure reason
### (now based on entry_clean since discard_date is separate)
# ------------------------------------------------------------

cells_analysis <- cells_analysis %>%
  mutate(
    failure_reason = case_when(
      str_detect(entry_clean,
        "fung|contam|bacteria")                      ~ "contamination",
      str_detect(entry_clean,
        "not growing|no growth|not gerowing|unhealthy|poor|slow|dying|failed") ~ "biological_failure",
      str_detect(entry_clean, "discarded")           ~ "discarded_unspecified",
      round_status == "complete"                     ~ NA_character_,
      round_status == "active"                       ~ NA_character_,
      TRUE                                           ~ NA_character_
    )
  )

# ------------------------------------------------------------
### STEP 15 — Final column selection
# ------------------------------------------------------------

cells_analysis <- cells_analysis %>%
  select(
    taxa, mv_parent, dish_id, common_name, species,
    tissue_type, dish_size, round_number, n_rounds_dish,
    entry_nature, round_status, failure_reason,
    date_collected, date_prep, discard_date, date_1st,
    p1_date, p2_date, p3_date, p4_date, p5_date, freeze_date,
    cell_n, pct_live, tube_n,
    days_collection_to_prep, days_prep_to_first,
    days_prep_to_discard, days_first_to_discard, days_p1_to_discard,
    days_first_to_p1, days_p1_to_p2,
    days_p2_to_freeze, days_prep_to_freeze,
    max_passage_reached, max_passage_label,
    outcome_A, outcome_B, outcome_C, outcome_D
  )

# ------------------------------------------------------------
### STEP 16 — Validation report
# ------------------------------------------------------------

cat("\n=== CVUI VALIDATION REPORT — Version 9 ===\n")
cat("Final n:", nrow(cells_analysis), "\n")
cat("Unique dish_id:", n_distinct(cells_analysis$dish_id), "\n")
cat("Unique species:", n_distinct(cells_analysis$species), "\n")

cat("\n--- n per taxa ---\n")
cells_analysis %>% count(taxa) %>% print()

cat("\n--- round_status distribution ---\n")
cells_analysis %>% count(round_status, sort = TRUE) %>% print()

cat("\n--- failure_reason distribution ---\n")
cells_analysis %>%
  count(failure_reason, sort = TRUE) %>%
  print(n = Inf)

cat("\n--- discard_date completeness ---\n")
cells_analysis %>%
  summarise(
    n_with_discard   = sum(!is.na(discard_date)),
    n_without_discard = sum(is.na(discard_date)),
    pct_complete     = round(100 * mean(!is.na(discard_date)), 1)
  ) %>%
  print()

cat("\n--- discard_date by outcome_A failure ---\n")
cells_analysis %>%
  filter(!is.na(outcome_A), outcome_A == 0L) %>%
  summarise(
    n_failures         = n(),
    n_with_discard     = sum(!is.na(discard_date)),
    n_without_discard  = sum(is.na(discard_date)),
    pct_complete       = round(100 * mean(!is.na(discard_date)), 1)
  ) %>%
  print()

cat("\n--- Predictor completeness (%) ---\n")
cells_analysis %>%
  summarise(across(
    c(cell_n, pct_live, days_collection_to_prep,
      days_prep_to_first, days_prep_to_discard,
      days_first_to_p1, days_p1_to_p2, days_p2_to_freeze),
    ~ round(mean(!is.na(.x)) * 100, 1)
  )) %>%
  pivot_longer(everything(),
               names_to  = "variable",
               values_to = "pct_complete") %>%
  print(n = Inf)

cat("\n--- Outcome distribution ---\n")
outcome_success_rates <- c(A = NA_real_, B = NA_real_, C = NA_real_, D = NA_real_)
for (oc in c("outcome_A", "outcome_B", "outcome_C", "outcome_D")) {
  n_elig <- sum(!is.na(cells_analysis[[oc]]))
  n_succ <- sum(cells_analysis[[oc]] == 1L, na.rm = TRUE)
  n_fail <- sum(cells_analysis[[oc]] == 0L, na.rm = TRUE)
  cat(sprintf(
    "  %s: eligible=%d, success=%d (%.0f%%), failure=%d (%.0f%%)\n",
    oc, n_elig, n_succ, 100 * n_succ / n_elig,
    n_fail, 100 * n_fail / n_elig
  ))
  letter <- toupper(sub("outcome_", "", oc))
  outcome_success_rates[letter] <- n_succ / n_elig
}

# ============================================================
### CVUI weight derivation — reproducible, data-driven
#
### Rationale: outcomes with lower observed success (higher
### attrition) represent harder biological gates and should
### contribute more to the total score. Each outcome's weight is
### set proportional to the reciprocal of its observed success
### rate (1 / success_rate), so weight increases as success rate
### falls. Reciprocals are scaled so the four stage weights sum
### to `total_points` (95 of 100; the remaining 5 points are
### reserved for the continuous Outcome D viability modifier).
#
### Raw scaled weights are not integers, so final weights are
### assigned via the largest-remainder (Hamilton apportionment)
### method: take the floor of each raw weight, then award the
### remaining points one at a time to the outcomes with the
### largest fractional remainders. This guarantees the weights
### are whole numbers that sum to exactly `total_points`,
### regardless of rounding.
# ============================================================

derive_cvui_weights <- function(success_rates, total_points = 95) {

  inv_rates   <- 1 / success_rates
  raw_weights <- inv_rates / sum(inv_rates) * total_points

  floor_weights <- floor(raw_weights)
  remainder     <- raw_weights - floor_weights
  deficit       <- total_points - sum(floor_weights)

  bump <- order(remainder, decreasing = TRUE)[seq_len(deficit)]
  floor_weights[bump] <- floor_weights[bump] + 1

  setNames(as.integer(floor_weights), names(success_rates))
}

cvui_weights <- derive_cvui_weights(outcome_success_rates, total_points = 95)

cat("\nDerived CVUI weights (reciprocal of success rate, scaled to 95, largest-remainder rounding):\n")
print(cvui_weights)

stopifnot(
  "Derived weights do not match the published CVUI V1 weights (A=29, B=28, C=18, D=20) — check data or derivation before proceeding" =
    all(cvui_weights == c(A = 29L, B = 28L, C = 18L, D = 20L))
)

cat("\n--- Outcome distribution by taxa ---\n")
cells_analysis %>%
  group_by(taxa) %>%
  summarise(
    n      = n(),
    n_A    = sum(!is.na(outcome_A)),
    prop_A = round(mean(outcome_A, na.rm = TRUE), 3),
    n_B    = sum(!is.na(outcome_B)),
    prop_B = round(mean(outcome_B, na.rm = TRUE), 3),
    n_C    = sum(!is.na(outcome_C)),
    prop_C = round(mean(outcome_C, na.rm = TRUE), 3),
    n_D    = sum(!is.na(outcome_D)),
    prop_D = round(mean(outcome_D, na.rm = TRUE), 3),
    .groups = "drop"
  ) %>%
  print()

cat("\n--- Negative interval check (all should be 0) ---\n")
cells_analysis %>%
  summarise(across(
    starts_with("days_"),
    ~ sum(.x < 0, na.rm = TRUE)
  )) %>%
  pivot_longer(everything(),
               names_to  = "variable",
               values_to = "n_negative") %>%
  print()

glimpse(cells_analysis)
```
#Survival Analysis - CVUI Outcome A only
```{r}
# ============================================================
### Survival analysis — Outcome A
### Event: failure to produce first cells (outcome_A == 0)
### Time: days_prep_to_discard for failures
#       days_prep_to_first for successes (censored)
### Random effect: mv_parent
# ============================================================

library(survival)
library(coxme)
library(dplyr)

cat("=== SURVIVAL ANALYSIS — OUTCOME A ===\n\n")

surv_data_A <- cells_analysis %>%
  filter(!is.na(outcome_A)) %>%
  mutate(
    # Time = days until event or censoring
    surv_time = case_when(
      outcome_A == 0L & !is.na(days_prep_to_discard) ~ days_prep_to_discard,
      outcome_A == 1L & !is.na(days_prep_to_first)   ~ days_prep_to_first,
      TRUE                                            ~ NA_real_
    ),
    # Event = failure (1) or censored/success (0)
    event_A = if_else(outcome_A == 0L, 1L, 0L),
    mv_parent = as.character(mv_parent)
  ) %>%
  filter(!is.na(surv_time), surv_time > 0)

cat("n for survival model:", nrow(surv_data_A), "\n")
cat("Events (failures — no cells):", sum(surv_data_A$event_A == 1L), "\n")
cat("Censored (cells appeared):", sum(surv_data_A$event_A == 0L), "\n")
cat("mv_parent clusters:", n_distinct(surv_data_A$mv_parent), "\n\n")

cat("--- Events vs censored by taxa ---\n")
surv_data_A %>%
  group_by(taxa) %>%
  summarise(
    n          = n(),
    n_events   = sum(event_A == 1L),
    n_censored = sum(event_A == 0L),
    .groups = "drop"
  ) %>%
  print()

cat("\n--- Kaplan-Meier by taxa ---\n")
km_A <- survfit(
  Surv(surv_time, event_A) ~ taxa,
  data = surv_data_A
)
print(km_A)

cat("\n--- Cox mixed effects: taxa + (1|mv_parent) ---\n")
cox_A <- coxme(
  Surv(surv_time, event_A) ~ taxa + (1 | mv_parent),
  data = surv_data_A
)
print(cox_A)

cat("\n--- Outcome B survival setup ---\n")
cat("Time = days_prep_to_first (present in all eligible)\n")
cat("Event = failure to reach P1 (outcome_B == 0)\n")
cat("Censored = reached P1 (outcome_B == 1)\n\n")

surv_data_B <- cells_analysis %>%
  filter(!is.na(outcome_B), !is.na(days_prep_to_first)) %>%
  mutate(
    surv_time = days_prep_to_first,
    event_B   = if_else(outcome_B == 0L, 1L, 0L),
    mv_parent = as.character(mv_parent)
  )

cat("n:", nrow(surv_data_B), "\n")
cat("Events (failures):", sum(surv_data_B$event_B == 1L), "\n")
cat("Censored (successes):", sum(surv_data_B$event_B == 0L), "\n\n")

cat("--- Cox mixed effects: taxa + (1|mv_parent) ---\n")
cox_B <- coxme(
  Surv(surv_time, event_B) ~ taxa + (1 | mv_parent),
  data = surv_data_B
)
print(cox_B)
```
#Survival analysis — Outcomes C and D
```{r}
# ============================================================
### Survival analysis — Outcomes C and D
# ============================================================

library(survival)
library(coxme)
library(dplyr)

# ------------------------------------------------------------
### OUTCOME C
### Event: failure to expand (outcome_C == 0)
### Time: days_first_to_p1 (present in all eligible — all passed B)
### Censored: reached P2 or freeze (outcome_C == 1)
# ------------------------------------------------------------

cat("=== SURVIVAL ANALYSIS — OUTCOME C ===\n\n")

surv_data_C <- cells_analysis %>%
  filter(!is.na(outcome_C), !is.na(days_first_to_p1)) %>%
  mutate(
    surv_time = days_first_to_p1,
    event_C   = if_else(outcome_C == 0L, 1L, 0L),
    mv_parent = as.character(mv_parent)
  )

cat("n:", nrow(surv_data_C), "\n")
cat("Events (failures):", sum(surv_data_C$event_C == 1L), "\n")
cat("Censored (successes):", sum(surv_data_C$event_C == 0L), "\n")
cat("mv_parent clusters:", n_distinct(surv_data_C$mv_parent), "\n\n")

cat("--- Events vs censored by taxa ---\n")
surv_data_C %>%
  group_by(taxa) %>%
  summarise(
    n          = n(),
    n_events   = sum(event_C == 1L),
    n_censored = sum(event_C == 0L),
    .groups = "drop"
  ) %>%
  print()

cat("\n--- Kaplan-Meier by taxa ---\n")
km_C <- survfit(
  Surv(surv_time, event_C) ~ taxa,
  data = surv_data_C
)
print(km_C)

cat("\n--- Cox mixed effects: taxa + (1|mv_parent) ---\n")
tryCatch({
  cox_C <- coxme(
    Surv(surv_time, event_C) ~ taxa + (1 | mv_parent),
    data = surv_data_C
  )
  print(cox_C)
}, error = function(e) {
  cat("Model error:", conditionMessage(e), "\n")
  cat("Trying fixed effects only:\n")
  cox_C_fixed <- coxph(
    Surv(surv_time, event_C) ~ taxa,
    data = surv_data_C
  )
  print(summary(cox_C_fixed))
})

# ------------------------------------------------------------
### OUTCOME D
### Event: failure to bank (outcome_D == 0)
### Time: days_p1_to_p2 (present in all eligible — all passed C)
### Censored: banked (outcome_D == 1)
# ------------------------------------------------------------

cat("\n\n=== SURVIVAL ANALYSIS — OUTCOME D ===\n\n")

surv_data_D <- cells_analysis %>%
  filter(!is.na(outcome_D), !is.na(days_p1_to_p2)) %>%
  mutate(
    surv_time = days_p1_to_p2,
    event_D   = if_else(outcome_D == 0L, 1L, 0L),
    mv_parent = as.character(mv_parent)
  )

cat("n:", nrow(surv_data_D), "\n")
cat("Events (failures):", sum(surv_data_D$event_D == 1L), "\n")
cat("Censored (successes):", sum(surv_data_D$event_D == 0L), "\n")
cat("mv_parent clusters:", n_distinct(surv_data_D$mv_parent), "\n\n")

cat("--- Events vs censored by taxa ---\n")
surv_data_D %>%
  group_by(taxa) %>%
  summarise(
    n          = n(),
    n_events   = sum(event_D == 1L),
    n_censored = sum(event_D == 0L),
    .groups = "drop"
  ) %>%
  print()

cat("\n--- Kaplan-Meier by taxa ---\n")
km_D <- survfit(
  Surv(surv_time, event_D) ~ taxa,
  data = surv_data_D
)
print(km_D)

cat("\n--- Cox mixed effects: taxa + (1|mv_parent) ---\n")
tryCatch({
  cox_D <- coxme(
    Surv(surv_time, event_D) ~ taxa + (1 | mv_parent),
    data = surv_data_D
  )
  print(cox_D)
}, error = function(e) {
  cat("Model error:", conditionMessage(e), "\n")
  cat("Trying fixed effects only:\n")
  cox_D_fixed <- coxph(
    Surv(surv_time, event_D) ~ taxa,
    data = surv_data_D
  )
  print(summary(cox_D_fixed))
})

# ------------------------------------------------------------
### Summary table across all outcomes
# ------------------------------------------------------------

cat("\n\n=== SURVIVAL ANALYSIS SUMMARY ===\n")
cat("Outcome A: n=66, events=10 (26% discard date complete) — underpowered\n")
cat("Outcome B: n=55, events=24, Mammal HR=3.618 p=0.009, overall p=0.039\n")
cat("Outcome C: see above\n")
cat("Outcome D: see above\n")
```
#CVUI Version 1 — Score calculation
```{r}

# ============================================================
### CVUI Version 1 — Score calculation
### Scale: 0–100
### Weights: derived above from observed success rates via
### derive_cvui_weights() — validated to equal A=29, B=28,
### C=18, D=20 (see stopifnot check)
### Quality modifier: continuous pct_live/100 × 5
# ============================================================

cells_analysis <- cells_analysis %>%
  mutate(

    # ------------------------------------------------------------
    # Stage scores (binary 0/1)
    # ------------------------------------------------------------

    score_A = case_when(
      outcome_A == 1L ~ 1L,
      outcome_A == 0L ~ 0L,
      TRUE            ~ NA_integer_
    ),

    score_B = case_when(
      is.na(outcome_B)  ~ 0L,  # did not reach B gate — counts as 0
      outcome_B == 1L   ~ 1L,
      outcome_B == 0L   ~ 0L,
      TRUE              ~ 0L
    ),

    score_C = case_when(
      is.na(outcome_C)  ~ 0L,
      outcome_C == 1L   ~ 1L,
      outcome_C == 0L   ~ 0L,
      TRUE              ~ 0L
    ),

    score_D = case_when(
      is.na(outcome_D)                              ~ 0L,
      outcome_D == 0L                               ~ 0L,
      # Failed freeze: 0% live AND no cell count
      outcome_D == 1L &
        !is.na(pct_live) & pct_live == 0 &
        is.na(cell_n)                               ~ 0L,
      outcome_D == 1L                               ~ 1L,
      TRUE                                          ~ 0L
    ),

    # ------------------------------------------------------------
    # Quality modifier at D — continuous pct_live/100
    # ------------------------------------------------------------

    quality_D = case_when(
      score_D == 0L                                 ~ 0,
      # Failed freeze quality
      !is.na(pct_live) & pct_live == 0 & is.na(cell_n) ~ 0,
      # Recorded viability — continuous score
      score_D == 1L & !is.na(pct_live) & pct_live > 0  ~ pct_live / 100,
      # Missing viability but freeze present — benefit of doubt
      score_D == 1L & is.na(pct_live)               ~ 1,
      TRUE                                          ~ 0
    ),

    # ------------------------------------------------------------
    # CVUI score — 0 to 100
    # ------------------------------------------------------------

    cvui_score = (score_A * cvui_weights["A"]) +
                 (score_B * cvui_weights["B"]) +
                 (score_C * cvui_weights["C"]) +
                 (score_D * cvui_weights["D"]) +
                 (quality_D * 5),

    # ------------------------------------------------------------
    # Flag for missing viability — for retrospective update
    # ------------------------------------------------------------

    viability_flag = case_when(
      score_D == 1L & is.na(pct_live) ~ "viability_missing",
      score_D == 1L & pct_live == 0 &
        is.na(cell_n)                 ~ "failed_freeze",
      score_D == 1L & pct_live > 0   ~ "viability_recorded",
      TRUE                            ~ NA_character_
    )
  )

# ------------------------------------------------------------
### Validation and summary
# ------------------------------------------------------------

cat("=== CVUI VERSION 1 SCORES ===\n\n")

cat("--- Score distribution ---\n")
cells_analysis %>%
  filter(!is.na(score_A)) %>%
  summarise(
    n         = n(),
    mean      = round(mean(cvui_score, na.rm = TRUE), 1),
    median    = round(median(cvui_score, na.rm = TRUE), 1),
    SD        = round(sd(cvui_score, na.rm = TRUE), 1),
    min       = min(cvui_score, na.rm = TRUE),
    max       = max(cvui_score, na.rm = TRUE)
  ) %>%
  print()

cat("\n--- Score distribution by taxa ---\n")
cells_analysis %>%
  filter(!is.na(score_A)) %>%
  group_by(taxa) %>%
  summarise(
    n      = n(),
    mean   = round(mean(cvui_score, na.rm = TRUE), 1),
    median = round(median(cvui_score, na.rm = TRUE), 1),
    SD     = round(sd(cvui_score, na.rm = TRUE), 1),
    min    = min(cvui_score, na.rm = TRUE),
    max    = max(cvui_score, na.rm = TRUE),
    .groups = "drop"
  ) %>%
  print()

cat("\n--- Score category distribution ---\n")
cells_analysis %>%
  filter(!is.na(score_A)) %>%
  mutate(
    cvui_category = case_when(
      cvui_score == 0        ~ "0 — no establishment",
      cvui_score == 29       ~ "29 — established only",
      cvui_score == 57       ~ "57 — reached P1",
      cvui_score == 75       ~ "75 — expanded, not banked",
      cvui_score >= 95       ~ "95-100 — banked",
      TRUE                   ~ "other"
    )
  ) %>%
  count(cvui_category, sort = TRUE) %>%
  print()

cat("\n--- Viability flag distribution ---\n")
cells_analysis %>%
  filter(score_D == 1L) %>%
  count(viability_flag) %>%
  print()

cat("\n--- Score distribution by max_passage_label ---\n")
cells_analysis %>%
  filter(!is.na(score_A)) %>%
  group_by(max_passage_label) %>%
  summarise(
    n          = n(),
    mean_cvui  = round(mean(cvui_score, na.rm = TRUE), 1),
    .groups = "drop"
  ) %>%
  arrange(desc(mean_cvui)) %>%
  print()
```
#Graphing CVUI - Figure 1
```{r}
# ============================================================
### CVUI — Grouped bar chart by score category and taxon
### Shown as proportions within each taxon (not raw counts),
### per co-author comment: taxon sample sizes are unequal
### (Bird n=59, Mammal n=34, Reptile n=13 eligible rounds), so
### raw counts would visually overweight Birds regardless of
### whether the underlying distribution shape actually differs.
### Each taxon's bars sum to 1.0 across the five score categories.
# ============================================================

library(ggplot2)
library(dplyr)
library(scales)

taxon_colours_light <- c(
  "Bird"    = "#85B7EB",
  "Mammal"  = "#5DCAA5",
  "Reptile" = "#F0997B"
)

plot_data <- cells_analysis %>%
  filter(!is.na(score_A)) %>%
  mutate(
    cvui_category = case_when(
      cvui_score == 0  ~ "0 — no establishment",
      cvui_score <= 29 ~ "1–29 — established",
      cvui_score <= 57 ~ "30–57 — reached P1",
      cvui_score <= 75 ~ "58–75 — expanded",
      cvui_score > 75  ~ "76–100 — banked"
    ),
    cvui_category = factor(cvui_category, levels = c(
      "0 — no establishment",
      "1–29 — established",
      "30–57 — reached P1",
      "58–75 — expanded",
      "76–100 — banked"
    ))
  ) %>%
  count(taxa, cvui_category, name = "n") %>%
  group_by(taxa) %>%
  mutate(
    taxa_total = sum(n),
    proportion = n / taxa_total
  ) %>%
  ungroup()

### Build legend labels that include each taxon's n, since the
### proportion bars on their own no longer show sample size
taxon_n <- plot_data %>%
  distinct(taxa, taxa_total) %>%
  mutate(label = paste0(taxa, " (n=", taxa_total, ")"))

taxon_labels <- setNames(taxon_n$label, taxon_n$taxa)

ggplot(plot_data, aes(x = cvui_category, y = proportion, fill = taxa)) +
  geom_bar(
    stat     = "identity",
    position = position_dodge(width = 0.75),
    width    = 0.65,
    colour   = NA
  ) +
  scale_fill_manual(values = taxon_colours_light, labels = taxon_labels) +
  scale_y_continuous(
    expand = expansion(mult = c(0, 0.05)),
    labels = percent_format(accuracy = 1)
  ) +
  labs(
    title    = "CVUI score category distribution by taxon",
    subtitle = "Proportion of each taxon's eligible culture rounds (n=106 total; Bird n=59, Mammal n=34, Reptile n=13)",
    x        = NULL,
    y        = "Proportion of rounds (within taxon)",
    fill     = NULL
  ) +
  theme_minimal(base_size = 9) +
  theme(
    legend.position    = "bottom",
    legend.key.size    = unit(0.4, "cm"),
    panel.grid.major.x = element_blank(),
    panel.grid.minor   = element_blank(),
    axis.text.x        = element_text(size = 10),
    strip.background   = element_blank(),
    plot.subtitle      = element_text(colour = "grey50", size = 10)
  )
```

#Graphing CVUI - Figure 2
```{r}
# ============================================================
### Figure 2 (redesign) — Stage-attrition curve by taxon
### Standard legend key instead of direct/repelled line labels
# ============================================================

library(ggplot2)
library(dplyr)
library(tidyr)

# ------------------------------------------------------------
### 1. Colours (locked, same as Figure 3)
# ------------------------------------------------------------

taxon_colours <- c(
  "Bird"    = "#378ADD",
  "Mammal"  = "#1D9E75",
  "Reptile" = "#D85A30"
)

# ------------------------------------------------------------
### 2. Compute proportion reaching each gate, per taxon
# ------------------------------------------------------------

stage_levels <- c("Attempted", "Established", "Reached P1", "Expanded", "Banked")

attrition_data <- cells_analysis %>%
  filter(!is.na(score_A)) %>%
  group_by(taxa) %>%
  summarise(
    n            = n(),
    Attempted    = 1,
    Established  = mean(cvui_score >= 29),
    `Reached P1` = mean(cvui_score >= 57),
    Expanded     = mean(cvui_score >= 75),
    Banked       = mean(cvui_score >= 95),
    .groups = "drop"
  ) %>%
  pivot_longer(
    cols      = all_of(stage_levels),
    names_to  = "stage",
    values_to = "proportion"
  ) %>%
  mutate(
    stage    = factor(stage, levels = stage_levels),
    stage_n  = as.numeric(stage),
    taxa_label = paste0(taxa, " (n=", n, ")"),
    taxa     = factor(taxa, levels = c("Bird", "Mammal", "Reptile"))
  )

### keep legend labels in the same taxa order, with n included
legend_labels <- attrition_data %>%
  distinct(taxa, taxa_label) %>%
  arrange(taxa) %>%
  pull(taxa_label)
names(legend_labels) <- attrition_data %>% distinct(taxa) %>% pull(taxa)

# ------------------------------------------------------------
### 3. Plot
# ------------------------------------------------------------

p2 <- ggplot(attrition_data, aes(x = stage_n, y = proportion,
                                  colour = taxa, group = taxa)) +

  # alternating stage bands for visual rhythm, drawn first
  annotate("rect", xmin = 1.5, xmax = 2.5, ymin = -Inf, ymax = Inf,
           fill = "grey96") +
  annotate("rect", xmin = 3.5, xmax = 4.5, ymin = -Inf, ymax = Inf,
           fill = "grey96") +

  # attrition steps — thin, dashed
  geom_step(direction = "hv", linewidth = 0.7, linetype = "dashed",
            lineend = "round") +

  # points: white-bordered filled circles
  geom_point(size = 4, shape = 21, stroke = 1.1,
             aes(fill = taxa), colour = "white") +

  scale_colour_manual(values = taxon_colours, labels = legend_labels) +
  scale_fill_manual(values = taxon_colours, labels = legend_labels) +
  scale_x_continuous(
    breaks = 1:5,
    labels = c("Attempted", "Established\n(\u226529)", "Reached P1\n(\u226557)",
               "Expanded\n(\u226575)", "Banked\n(\u226595)"),
    limits = c(0.5, 5.1),
    expand = c(0, 0)
  ) +
  scale_y_continuous(
    labels = scales::percent,
    limits = c(0, 1),
    breaks = seq(0, 1, 0.25),
    expand = expansion(mult = c(0.02, 0.05))
  ) +
  labs(x = NULL, y = "Proportion of rounds reaching stage",
       colour = NULL, fill = NULL) +
  theme_minimal(base_size = 17) +
  theme(
    legend.position     = "top",
    legend.text         = element_text(size = 13),
    axis.text.x         = element_text(size = 14, colour = "grey20", lineheight = 1.1),
    axis.text.y         = element_text(size = 13, colour = "grey30"),
    axis.title.y        = element_text(size = 15, margin = margin(r = 12)),
    panel.grid.major.x  = element_blank(),
    panel.grid.minor    = element_blank(),
    panel.grid.major.y  = element_line(colour = "grey88", linewidth = 0.4),
    plot.margin         = margin(t = 15, r = 20, b = 15, l = 15)
  )

print(p2)
```
#Graphing CVUI - Figure 3
```{r}
# ============================================================
### Figure 3 — Violin + boxplot + jitter, no axis titles
# ============================================================

library(ggplot2)
library(dplyr)

# ------------------------------------------------------------
### 1. Colours (locked)
# ------------------------------------------------------------

taxon_colours <- c(
  "Bird"    = "#378ADD",
  "Mammal"  = "#1D9E75",
  "Reptile" = "#D85A30"
)

# ------------------------------------------------------------
### 2. Data prep
# ------------------------------------------------------------

plot_data <- cells_analysis %>%
  filter(!is.na(score_A)) %>%
  mutate(taxa = factor(taxa, levels = c("Bird", "Mammal", "Reptile")))

# ------------------------------------------------------------
### 3. Plot
# ------------------------------------------------------------

p3 <- ggplot(plot_data, aes(x = taxa, y = cvui_score, fill = taxa)) +

  geom_violin(trim = TRUE, alpha = 0.85, colour = "grey30", linewidth = 0.4) +

  geom_boxplot(width = 0.11, fill = "whitesmoke", outlier.shape = NA,
               colour = "grey20", linewidth = 0.5) +

  geom_jitter(width = 0.08, height = 0, size = 1.8, colour = "black", alpha = 0.9) +

  scale_fill_manual(values = taxon_colours, name = "Taxon") +
  scale_y_continuous(breaks = c(0, 29, 57, 75, 95, 100)) +
  labs(x = NULL, y = NULL) +
  theme_gray(base_size = 14)

print(p3)
```

#============================================================
# Supplementary regression diagnostics — Outcomes A–D
# Integrated from the original regression script so that every
# block below runs against the SAME cells_analysis object built
# once at the top of this file (V9/V10 pipeline, with
# discard_date). This removes the earlier inconsistency where
# eligible/model n's differed across separate sessions.
#
# Note: the original script contained a broken, truncated
# duplicate of Block 3 (it cut off mid-expression) immediately
# before the working version. That duplicate has been removed
# here; only the complete, functional Block 3 is included below.
#============================================================

#Outcome A & B — descriptive and univariate regression diagnostics
```{r}
# ============================================================
### Outcome A descriptive + Outcome B univariate logistic regression
### Outcome A — descriptive only: days_prep_to_first is structurally
#             missing for failures (no cells ever appeared), so no
#             regression is fit; only descriptive success rates are
#             reported.
### Outcome B — predictor: days_prep_to_first
#             (present in both successes and failures)
# ============================================================

library(dplyr)
library(tidyr)
library(purrr)
library(pROC)

# ------------------------------------------------------------
### BLOCK 1 — Outcome A descriptive only
# ------------------------------------------------------------

cat("=== OUTCOME A — DESCRIPTIVE ONLY ===\n\n")

cat("--- Overall success rate ---\n")
cells_analysis %>%
  filter(!is.na(outcome_A)) %>%
  summarise(
    n_eligible  = n(),
    n_success   = sum(outcome_A == 1L),
    n_failure   = sum(outcome_A == 0L),
    pct_success = round(mean(outcome_A) * 100, 1)
  ) %>%
  print()

cat("\n--- Success rate by taxa ---\n")
cells_analysis %>%
  filter(!is.na(outcome_A)) %>%
  group_by(taxa) %>%
  summarise(
    n_eligible  = n(),
    n_success   = sum(outcome_A == 1L),
    n_failure   = sum(outcome_A == 0L),
    pct_success = round(mean(outcome_A) * 100, 1),
    .groups = "drop"
  ) %>%
  print()

cat("\n--- days_prep_to_first in SUCCESS group only ---\n")
cat("(Structurally missing in failures)\n\n")
cells_analysis %>%
  filter(!is.na(outcome_A), outcome_A == 1L,
         !is.na(days_prep_to_first)) %>%
  group_by(taxa) %>%
  summarise(
    n      = n(),
    mean   = round(mean(days_prep_to_first), 2),
    median = round(median(days_prep_to_first), 2),
    SD     = round(sd(days_prep_to_first), 2),
    .groups = "drop"
  ) %>%
  print()

# ------------------------------------------------------------
### BLOCK 2 — Outcome B regression
### Predictor: days_prep_to_first
### All Outcome B eligible have date_1st so interval present
### in both successes and failures
# ------------------------------------------------------------

cat("\n\n=== UNIVARIATE LOGISTIC REGRESSION — OUTCOME B ===\n")
cat("Predictor: days_prep_to_first\n\n")

model_data_B <- cells_analysis %>%
  filter(!is.na(outcome_B), !is.na(days_prep_to_first))

cat("n for model:", nrow(model_data_B), "\n")
cat("Failures:", sum(model_data_B$outcome_B == 0L), "\n")
cat("Successes:", sum(model_data_B$outcome_B == 1L), "\n\n")

if (nrow(model_data_B) > 5 &&
    sum(model_data_B$outcome_B == 0L) >= 3 &&
    sum(model_data_B$outcome_B == 1L) >= 3) {

  mod_B <- glm(
    outcome_B ~ days_prep_to_first,
    data   = model_data_B,
    family = binomial(link = "logit")
  )

  cat("--- Model summary ---\n")
  print(summary(mod_B))

  cat("\n--- Odds ratio and 95% CI ---\n")
  or_B <- data.frame(
    predictor = "days_prep_to_first",
    OR        = round(exp(coef(mod_B)["days_prep_to_first"]), 3),
    CI_lower  = round(exp(confint(mod_B)["days_prep_to_first", 1]), 3),
    CI_upper  = round(exp(confint(mod_B)["days_prep_to_first", 2]), 3),
    p_value   = round(summary(mod_B)$coefficients[
      "days_prep_to_first", 4], 4)
  )
  print(or_B)

  pred_B <- predict(mod_B, type = "response")
  roc_B  <- roc(model_data_B$outcome_B, pred_B, quiet = TRUE)
  cat("\nAUC:", round(auc(roc_B), 3), "\n")
  cat("AUC 95% CI:", round(ci.auc(roc_B)[1], 3), "-",
      round(ci.auc(roc_B)[3], 3), "\n")

} else {
  cat("Insufficient data for Outcome B model\n")
}

cat("\n--- Outcome B descriptive: days_prep_to_first ---\n")
model_data_B %>%
  mutate(outcome_group = if_else(outcome_B == 1L, "Success", "Failure")) %>%
  group_by(outcome_group) %>%
  summarise(
    n      = n(),
    mean   = round(mean(days_prep_to_first), 2),
    median = round(median(days_prep_to_first), 2),
    SD     = round(sd(days_prep_to_first), 2),
    .groups = "drop"
  ) %>%
  print()

cat("\n--- Outcome B taxa-stratified ---\n")
for (tx in c("Bird", "Mammal", "Reptile")) {
  tx_data <- cells_analysis %>%
    filter(taxa == tx,
           !is.na(outcome_B),
           !is.na(days_prep_to_first))

  cat("\n  Taxa:", tx,
      "| n:", nrow(tx_data),
      "| Failures:", sum(tx_data$outcome_B == 0L),
      "| Successes:", sum(tx_data$outcome_B == 1L), "\n")

  if (sum(tx_data$outcome_B == 0L) < 3 ||
      sum(tx_data$outcome_B == 1L) < 3) {
    cat("  Insufficient n — skipping\n")
    next
  }

  tryCatch({
    mod_tx <- glm(
      outcome_B ~ days_prep_to_first,
      data   = tx_data,
      family = binomial(link = "logit")
    )
    or_tx <- data.frame(
      taxa     = tx,
      OR       = round(exp(coef(mod_tx)["days_prep_to_first"]), 3),
      CI_lower = round(exp(confint(mod_tx)["days_prep_to_first", 1]), 3),
      CI_upper = round(exp(confint(mod_tx)["days_prep_to_first", 2]), 3),
      p_value  = round(summary(mod_tx)$coefficients[
        "days_prep_to_first", 4], 4)
    )
    print(or_tx)
    pred_tx <- predict(mod_tx, type = "response")
    roc_tx  <- roc(tx_data$outcome_B, pred_tx, quiet = TRUE)
    cat("  AUC:", round(auc(roc_tx), 3), "\n")
  }, error = function(e) {
    cat("  Model error:", conditionMessage(e), "\n")
  })
}
```

#Outcome C & D — univariate regression diagnostics
```{r}
# ------------------------------------------------------------
### BLOCK 3 — Outcome C regression
### Predictor: days_first_to_p1
### All Outcome C eligible have p1_date so interval present
### in both successes and failures
# ------------------------------------------------------------

cat("\n\n=== UNIVARIATE LOGISTIC REGRESSION — OUTCOME C ===\n")
cat("Predictor: days_first_to_p1\n\n")

model_data_C <- cells_analysis %>%
  filter(!is.na(outcome_C), !is.na(days_first_to_p1))

cat("n for model:", nrow(model_data_C), "\n")
cat("Failures:", sum(model_data_C$outcome_C == 0L), "\n")
cat("Successes:", sum(model_data_C$outcome_C == 1L), "\n\n")

if (nrow(model_data_C) > 5 &&
    sum(model_data_C$outcome_C == 0L) >= 3 &&
    sum(model_data_C$outcome_C == 1L) >= 3) {

  mod_C <- glm(
    outcome_C ~ days_first_to_p1,
    data   = model_data_C,
    family = binomial(link = "logit")
  )

  cat("--- Model summary ---\n")
  print(summary(mod_C))

  cat("\n--- Odds ratio and 95% CI ---\n")
  or_C <- data.frame(
    predictor = "days_first_to_p1",
    OR        = round(exp(coef(mod_C)["days_first_to_p1"]), 3),
    CI_lower  = round(exp(confint(mod_C)["days_first_to_p1", 1]), 3),
    CI_upper  = round(exp(confint(mod_C)["days_first_to_p1", 2]), 3),
    p_value   = round(summary(mod_C)$coefficients[
      "days_first_to_p1", 4], 4)
  )
  print(or_C)

  pred_C <- predict(mod_C, type = "response")
  roc_C  <- roc(model_data_C$outcome_C, pred_C, quiet = TRUE)
  cat("\nAUC:", round(auc(roc_C), 3), "\n")
  cat("AUC 95% CI:", round(ci.auc(roc_C)[1], 3), "-",
      round(ci.auc(roc_C)[3], 3), "\n")

} else {
  cat("Insufficient data for Outcome C model\n")
}

cat("\n--- Outcome C descriptive: days_first_to_p1 ---\n")
model_data_C %>%
  mutate(outcome_group = if_else(outcome_C == 1L, "Success", "Failure")) %>%
  group_by(outcome_group) %>%
  summarise(
    n      = n(),
    mean   = round(mean(days_first_to_p1), 2),
    median = round(median(days_first_to_p1), 2),
    SD     = round(sd(days_first_to_p1), 2),
    .groups = "drop"
  ) %>%
  print()

# ------------------------------------------------------------
### BLOCK 4 — Outcome D regression
### Predictor: days_p1_to_p2
# ------------------------------------------------------------

cat("\n\n=== UNIVARIATE LOGISTIC REGRESSION — OUTCOME D ===\n")
cat("Predictor: days_p1_to_p2\n\n")

model_data_D <- cells_analysis %>%
  filter(!is.na(outcome_D), !is.na(days_p1_to_p2))

cat("n for model:", nrow(model_data_D), "\n")
cat("Failures:", sum(model_data_D$outcome_D == 0L), "\n")
cat("Successes:", sum(model_data_D$outcome_D == 1L), "\n\n")

if (nrow(model_data_D) > 5 &&
    sum(model_data_D$outcome_D == 0L) >= 3 &&
    sum(model_data_D$outcome_D == 1L) >= 3) {

  mod_D <- glm(
    outcome_D ~ days_p1_to_p2,
    data   = model_data_D,
    family = binomial(link = "logit")
  )

  cat("--- Model summary ---\n")
  print(summary(mod_D))

  cat("\n--- Odds ratio and 95% CI ---\n")
  or_D <- data.frame(
    predictor = "days_p1_to_p2",
    OR        = round(exp(coef(mod_D)["days_p1_to_p2"]), 3),
    CI_lower  = round(exp(confint(mod_D)["days_p1_to_p2", 1]), 3),
    CI_upper  = round(exp(confint(mod_D)["days_p1_to_p2", 2]), 3),
    p_value   = round(summary(mod_D)$coefficients[
      "days_p1_to_p2", 4], 4)
  )
  print(or_D)

  pred_D <- predict(mod_D, type = "response")
  roc_D  <- roc(model_data_D$outcome_D, pred_D, quiet = TRUE)
  cat("\nAUC:", round(auc(roc_D), 3), "\n")
  cat("AUC 95% CI:", round(ci.auc(roc_D)[1], 3), "-",
      round(ci.auc(roc_D)[3], 3), "\n")

} else {
  cat("Insufficient data for Outcome D model\n")
}

cat("\n--- Outcome D descriptive: days_p1_to_p2 ---\n")
model_data_D %>%
  mutate(outcome_group = if_else(outcome_D == 1L, "Success", "Failure")) %>%
  group_by(outcome_group) %>%
  summarise(
    n      = n(),
    mean   = round(mean(days_p1_to_p2), 2),
    median = round(median(days_p1_to_p2), 2),
    SD     = round(sd(days_p1_to_p2), 2),
    .groups = "drop"
  ) %>%
  print()
```

#Testing possible multi-regression: see how many levels tissue type has in your Outcome B eligible group.
```{r}
### Check tissue type distribution in Outcome B eligible rounds
cells_analysis %>%
  filter(!is.na(outcome_B)) %>%
  count(tissue_type, outcome_B, sort = TRUE) %>%
  print(n = Inf)

### Check taxa distribution
cells_analysis %>%
  filter(!is.na(outcome_B)) %>%
  count(taxa, outcome_B) %>%
  print(n = Inf)
```

#checking mv_parent clustering
```{r}
# ============================================================
### Check clustering structure for mixed effects model
### Unit: mv_parent as random effect
# ============================================================

cat("=== CLUSTERING STRUCTURE — mv_parent ===\n\n")

cat("--- Total unique mv_parent values ---\n")
cat(n_distinct(cells_analysis$mv_parent), "\n\n")

cat("--- Distribution of observations per mv_parent ---\n")
cells_analysis %>%
  count(mv_parent, name = "n_obs") %>%
  count(n_obs, name = "n_parents") %>%
  mutate(pct = round(100 * n_parents / sum(n_parents), 1)) %>%
  print(n = Inf)

cat("\n--- mv_parent with multiple observations ---\n")
cells_analysis %>%
  count(mv_parent, name = "n_obs") %>%
  filter(n_obs > 1) %>%
  arrange(desc(n_obs)) %>%
  print(n = Inf)

cat("\n--- Clustering within Outcome B eligible ---\n")
cells_analysis %>%
  filter(!is.na(outcome_B)) %>%
  count(mv_parent, name = "n_obs") %>%
  count(n_obs, name = "n_parents") %>%
  print(n = Inf)

cat("\n--- How many mv_parent have both successes and failures at Outcome B ---\n")
cells_analysis %>%
  filter(!is.na(outcome_B)) %>%
  group_by(mv_parent) %>%
  summarise(
    n_obs      = n(),
    n_success  = sum(outcome_B == 1L),
    n_failure  = sum(outcome_B == 0L),
    .groups = "drop"
  ) %>%
  filter(n_obs > 1) %>%
  print(n = Inf)
```

#Mixed effects logistic regression — Outcome B (random intercept for mv_parent; no taxa term)
```{r}
# ============================================================
### Mixed effects logistic regression — Outcome B
### Random intercept for mv_parent
### (NOTE: corrected header above — the original comment said
### "with bird as the reference taxa," but this model has no
### taxa term at all; that mismatch was in the original script,
### not introduced here.)
# ============================================================

library(lme4)
library(pROC)

cat("=== MIXED EFFECTS MODEL — OUTCOME B ===\n")
cat("Fixed: days_prep_to_first\n")
cat("Random: (1 | mv_parent)\n\n")

### NOTE: explicitly excluding rows with missing mv_parent. Without this,
### glmer() silently drops them internally (since the grouping variable
### is NA), so its "Number of obs" ends up one less than nrow() of the
### input data — and predict()/roc() then fail on a length mismatch
### between predictions and the full outcome vector. Filtering here keeps
### the printed n, the model's internal n, and the AUC calculation all
### referring to the same set of rows.
model_data_B_mixed <- cells_analysis %>%
  filter(!is.na(outcome_B), !is.na(days_prep_to_first), !is.na(mv_parent)) %>%
  mutate(mv_parent = as.factor(mv_parent))

cat("n observations:", nrow(model_data_B_mixed), "\n")
cat("n mv_parent clusters:", n_distinct(model_data_B_mixed$mv_parent), "\n")
cat("Failures:", sum(model_data_B_mixed$outcome_B == 0L), "\n")
cat("Successes:", sum(model_data_B_mixed$outcome_B == 1L), "\n\n")

mod_B_mixed <- glmer(
  outcome_B ~ days_prep_to_first + (1 | mv_parent),
  data   = model_data_B_mixed,
  family = binomial(link = "logit"),
  control = glmerControl(optimizer = "bobyqa",
                         optCtrl = list(maxfun = 2e5))
)

cat("--- Model summary ---\n")
print(summary(mod_B_mixed))

cat("\n--- Fixed effects OR and 95% CI ---\n")
or_mixed <- data.frame(
  predictor = "days_prep_to_first",
  OR        = round(exp(fixef(mod_B_mixed)["days_prep_to_first"]), 3),
  CI_lower  = round(exp(confint(mod_B_mixed,
    parm = "days_prep_to_first", method = "Wald")[1]), 3),
  CI_upper  = round(exp(confint(mod_B_mixed,
    parm = "days_prep_to_first", method = "Wald")[2]), 3),
  p_value   = round(summary(mod_B_mixed)$coefficients[
    "days_prep_to_first", 4], 4)
)
print(or_mixed)

cat("\n--- Random effect variance (mv_parent) ---\n")
print(VarCorr(mod_B_mixed))

cat("\n--- ICC (intraclass correlation) ---\n")
icc_val <- as.numeric(VarCorr(mod_B_mixed)$mv_parent) /
  (as.numeric(VarCorr(mod_B_mixed)$mv_parent) + (pi^2 / 3))
cat("ICC:", round(icc_val, 3), "\n")
cat("(proportion of variance attributable to mv_parent)\n")

cat("\n--- AUC ---\n")
### NOTE: re.form = NA forces population-level predictions (random
### effects set to zero), not subject-specific ones. Without this,
### predict() uses each row's own fitted random intercept — and since
### 24 of the 33 mv_parent clusters here have only one observation,
### glmer can fit an intercept that drives that single prediction
### arbitrarily close to the observed outcome. That produces an AUC
### inflated by the model partly "predicting" an animal's outcome
### using information from that same animal's outcome (leakage), not
### genuine discrimination from the fixed effect alone. re.form = NA
### gives the AUC that's actually comparable to the fixed-effects-only
### model above.
pred_mixed <- predict(mod_B_mixed, type = "response", re.form = NA)
roc_mixed  <- roc(model_data_B_mixed$outcome_B, pred_mixed, quiet = TRUE)
cat("AUC:", round(auc(roc_mixed), 3), "\n")

cat("\n--- Compare fixed vs mixed model AIC ---\n")
cat("Fixed only AIC:", round(AIC(mod_B), 3), "\n")
cat("Mixed model AIC:", round(AIC(mod_B_mixed), 3), "\n")
```

#Taxa comparison — Outcome B, Mammal as reference (Reptile vs Mammal, Bird vs Mammal)
```{r}
# ------------------------------------------------------------
### Rerun with Mammal as reference to get Reptile vs Mammal
### and Bird vs Mammal directly
# ------------------------------------------------------------

model_data_B_reptile_ref <- cells_analysis %>%
  filter(!is.na(outcome_B), !is.na(days_prep_to_first)) %>%
  mutate(taxa = factor(taxa, levels = c("Mammal", "Bird", "Reptile")))

mod_B_mammal_ref <- glm(
  outcome_B ~ days_prep_to_first + taxa,
  data   = model_data_B_reptile_ref,
  family = binomial(link = "logit")
)

cat("--- Mammal as reference — Bird vs Mammal, Reptile vs Mammal ---\n")
or_mammal_ref <- data.frame(
  predictor = names(coef(mod_B_mammal_ref))[-1],
  OR        = round(exp(coef(mod_B_mammal_ref))[-1], 3),
  CI_lower  = round(exp(confint(mod_B_mammal_ref))[-1, 1], 3),
  CI_upper  = round(exp(confint(mod_B_mammal_ref))[-1, 2], 3),
  p_value   = round(summary(mod_B_mammal_ref)$coefficients[-1, 4], 4)
)
print(or_mammal_ref)
```

#Mixed effects model — Outcome B with taxa (Mammal as reference, random intercept for mv_parent)
```{r}
# ============================================================
### Mixed effects logistic regression — Outcome B, with taxa
### Fixed effects: days_prep_to_first + taxa (Mammal = reference,
#   consistent with the flat model above)
### Random intercept: mv_parent
#
### This is the model the earlier mislabeled chunk header ("with
### bird as the reference taxa") was presumably meant to describe
### — a taxa term was never actually fitted alongside the random
### intercept in that chunk. Built here properly instead of just
### relabeling the comment. Reference level kept as Mammal, not
### Bird, to stay consistent with the flat model immediately above
### rather than introduce a third, inconsistent reference scheme.
#
### This directly parallels the Cox mixed-effects model already
### fitted for Outcome B (taxon fixed effect + animal random
### intercept), but on the logistic/AUC side of the analysis, and
### accounts for repeated culture rounds from the same individual
### animal in a way the flat Mammal-reference model above does not.
# ============================================================

library(lme4)
library(pROC)

model_data_B_taxa_mixed <- cells_analysis %>%
  filter(!is.na(outcome_B), !is.na(days_prep_to_first), !is.na(mv_parent)) %>%
  mutate(
    mv_parent = as.factor(mv_parent),
    taxa      = factor(taxa, levels = c("Mammal", "Bird", "Reptile"))
  )

cat("=== MIXED EFFECTS MODEL — OUTCOME B (WITH TAXA, MAMMAL REFERENCE) ===\n")
cat("Fixed: days_prep_to_first + taxa\n")
cat("Random: (1 | mv_parent)\n\n")

cat("n observations:", nrow(model_data_B_taxa_mixed), "\n")
cat("n mv_parent clusters:", n_distinct(model_data_B_taxa_mixed$mv_parent), "\n\n")

mod_B_taxa_mixed <- glmer(
  outcome_B ~ days_prep_to_first + taxa + (1 | mv_parent),
  data    = model_data_B_taxa_mixed,
  family  = binomial(link = "logit"),
  control = glmerControl(optimizer = "bobyqa",
                         optCtrl = list(maxfun = 2e5))
)

cat("--- Model summary ---\n")
print(summary(mod_B_taxa_mixed))

cat("\n--- Fixed effects OR and 95% CI (Wald) ---\n")
fixed_terms   <- names(fixef(mod_B_taxa_mixed))[-1]  # drop intercept
ci_taxa_mixed <- confint(mod_B_taxa_mixed, parm = fixed_terms, method = "Wald")

or_taxa_mixed <- data.frame(
  predictor = fixed_terms,
  OR        = round(exp(fixef(mod_B_taxa_mixed)[fixed_terms]), 3),
  CI_lower  = round(exp(ci_taxa_mixed[, 1]), 3),
  CI_upper  = round(exp(ci_taxa_mixed[, 2]), 3),
  p_value   = round(summary(mod_B_taxa_mixed)$coefficients[fixed_terms, 4], 4)
)
print(or_taxa_mixed)

cat("\n--- Random effect variance (mv_parent) ---\n")
print(VarCorr(mod_B_taxa_mixed))

cat("\n--- ICC (intraclass correlation) ---\n")
icc_taxa_mixed <- as.numeric(VarCorr(mod_B_taxa_mixed)$mv_parent) /
  (as.numeric(VarCorr(mod_B_taxa_mixed)$mv_parent) + (pi^2 / 3))
cat("ICC:", round(icc_taxa_mixed, 3), "\n")

cat("\n--- Compare to flat (Mammal-ref, no random effect) model ---\n")
cat("Flat model AIC: ", round(AIC(mod_B_mammal_ref), 3), "\n")
cat("Mixed model AIC:", round(AIC(mod_B_taxa_mixed), 3), "\n")
cat("(Descriptive AIC comparison only — not a formal likelihood-ratio test)\n")
```
````
